# A Germinal Center Checkpoint of AIRE in B Cells Limits Antibody Diversification

**DOI:** 10.1101/2024.01.10.574926

**Authors:** Jordan Z. Zhou, Bo Pei, Guang Wen Sun, Michael D. Pawlitz, Wei Zhang, Xinyang Li, Fayi Yao, Kati C. Hokynar, Chun-Xiao Li, Madusha L. W. Perera, Shanqiao Wei, Simin Zheng, Lisa A. Polin, Janet M. Poulik, Annamari Ranki, Kai Krohn, Charlotte Cunningham-Rundles, Andrea Cerutti, Naibo Yang, Ashok S. Bhagwat, Kefei Yu, Pärt Peterson, Kai Kisand, Bao Q. Vuong, Bihui Huang, Kang Chen

**Author notes:** Correspondence (J.Z.Z.), (B.H.) and (K.C.). The authors dedicate this work to the memory of Dr. Annamari Ranki and Dr. Kai Krohn.

## Abstract

In response to antigens, B cells undergo antibody diversification, including affinity maturation and class switching, mediated by activation-induced cytidine deaminase (AID) in secondary lymphoid organs, but uncontrolled AID activity can precipitate autoimmunity and cancer. The regulation of antibody diversification is of fundamental importance, though the mechanisms underlying these pathways are not well understood. We found that autoimmune regulator (AIRE), the molecule essential for T cell tolerance, is expressed in germinal center (GC) B cells in a CD40-dependent manner, interacts with AID and negatively regulates antibody affinity maturation and class switching by inhibiting AID function. AIRE deficiency in B cells caused altered antibody repertoire, increased somatic hypermutations, elevated autoantibodies to T helper 17 effector cytokines and defective control of skin *Candida albicans*. These results define a GC B cell checkpoint of humoral immunity and illuminate new approaches of generating high-affinity neutralizing antibodies for immunotherapy.

## INTRODUCTION

A properly diversified and selected repertoire of antibodies is essential for effective immune defense against diverse pathogens as well as the prevention of autoimmune diseases. After V(D)J recombination in the bone marrow (BM) that generates a primary antibody repertoire, B lymphocytes enter secondary lymphoid organs, such as the lymph nodes, spleen, tonsils and mucosa-associated lymphoid tissues, and upon antigen stimulation, further diversify their antibody repertoire via class switch recombination (CSR) and somatic hypermutation (SHM), both of which are mediated by the DNA-cytosine deaminase AID (Muramatsu et al., 2000; Revy et al., 2000). SHM introduces point mutations in immunoglobulin (Ig) variable region exons for selection of higher-affinity antibody clones by antigens, whereas CSR replaces Ig constant region exons encoding IgM and IgD with those encoding IgG, IgA, or IgE to provide antibodies with new effector functions (Murphy and Weaver, 2016b). However, aberrant AID activity in B cells can cause mutations in non-Ig loci to precipitate cancer (Casellas et al., 2016), and AID-mediated germinal center (GC) reaction can generate autoreactive antibodies to drive many autoimmune diseases (Vinuesa et al., 2009). The absence of B cell lymphomas and overt autoimmune diseases in healthy individuals amidst ongoing humoral immune responses reflects the existence of physiological mechanisms that restrain AID-mediated antibody diversification in GC B cells. The details of these mechanisms are of fundamental importance but are not fully understood.

Humoral immunity is closely regulated by cell-mediated immunity, in which the molecule AIRE induces the expression of peripheral tissue-specific antigens (TSAs) in medullary epithelial cells (mTECs) and B cells in the thymus to promote the negative selection of self-reactive T cells or their conversion into regulatory T (Treg) cells (Anderson et al., 2002; Malchow et al., 2013; Yamano et al., 2015). AIRE is also expressed in specialized extrathymic cells (eTACs) that can inactivate self-reactive T cells in the periphery (Gardner et al., 2008; Gardner et al., 2013). Loss-of-function mutations in the *AIRE* gene cause autoimmune polyglandular syndrome type 1 (APS-1) (Finnish-German, 1997; Nagamine et al., 1997) associated with organ-specific autoimmunity, aberrant production of autoantibodies and increased susceptibility to mucocutaneous infection by *Candida albicans*, an otherwise innocuous commensal microbe in humans. Mysteriously, APS-1 patients can produce high-affinity neutralizing antibodies against T helper 17 (T_H_17) effector cytokines, which has been suggested to negatively impact anti-fungal immune defense (Kisand et al., 2010; Meyer et al., 2016a; Puel et al., 2010). In light of these findings, we sought to determine whether AIRE has a B cell-intrinsic role in regulating peripheral antibody diversification.

Here we show that AIRE is expressed in GC B cells in a CD40-dependent manner, interacts with AID, and negatively regulates AID-mediated peripheral antibody diversification. AIRE-deficient mouse B cells undergo elevated class switching and affinity maturation after antigenic stimulation, which correlates with enhanced generation of genomic uracil, elevated Ig SHM, augmented AID targeting to Ig switch (S) regions and increased interaction of AID with transcriptionally stalled RNA polymerase II (Pol II). In addition, naive B cells of APS-1 patients undergo increased CSR upon stimulation. Mice with AIRE deficiency in B cells have elevated levels of autoantibodies against T helper 17 (T_H_17) effector cytokines and heightened skin *C. albicans* burden after infection, which recapitulates the symptoms of APS-1 patients. Our results define a previously unknown but crucial B cell-intrinsic AIRE-mediated GC checkpoint of peripheral antibody diversification that limits autoimmunity.

## RESULTS

### GC B Cells Express AIRE

Secondary lymphoid organs are the major sites for peripheral antibody diversification (Murphy and Weaver, 2016a). To determine any roles of AIRE in peripheral antibody diversification, we first examined AIRE expression in B cells of human secondary lymphoid organs using an antibody that detects AIRE in the nuclei of mTECs (**Figure S1A**). We found many IgD^−^CD19^+^ or IgD^−^Pax5^+^ B cells inside tonsillar and splenic follicles that harbored nuclear AIRE (**Figure 1A–D**, **Figure S1B** and **C**). Follicular AIRE^+^ B cells expressed the GC B cell-associated molecule Bcl-6 (**Figure S1C**). In contrast, tonsillar and splenic IgD^+^ B cells in the mantle zone and IgD^+^ plasmablasts in GCs and extrafollicular areas (Chen et al., 2009) expressed little or no AIRE (**Figure S1B** and **Figure 1D**). Peripheral blood IgD^+^CD27^−^ or CD24^+^CD38^lo^ naive, IgD^+^CD27^+^ circulating marginal zone, IgD^−^CD27^+^ or CD24^hi^CD38^−^ memory, IgD^−^CD27^−^ atypical memory and CD24^hi^CD38^hi^ transitional B cells as well as CD24^−^CD38^hi^ plasma cells (PCs) did not express AIRE either (**Figure S1D**). Consistent with their follicular localization, tonsillar AIRE^+^ B cells were mostly IgD^−^CD38^+^ GC B cells, with a small fraction being IgD^+^CD38^+^ founder GC (FGC) or IgD^−^CD38^−^ memory B cells (**Figure 1E**).

**Figure 1.**
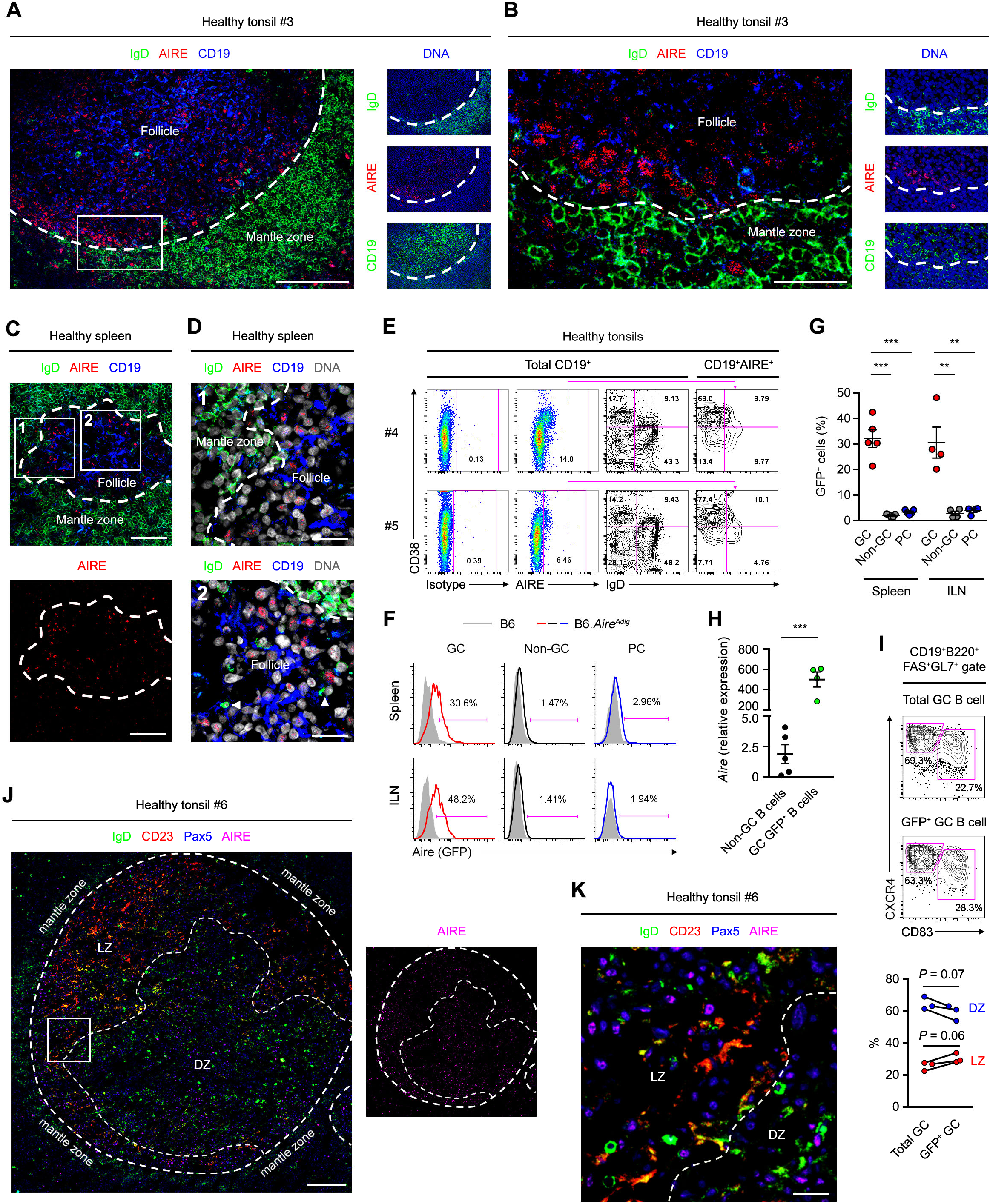
GC B Cells Express AIRE. (A and B) Immunofluorescence analysis of the tonsillar tissue of a healthy donor for IgD, CD19, AIRE and DAPI-stained DNA. The dotted line outlines the follicles. Bars: 100 μm (A) or 25 μm (B). (C and D) Immunofluorescence analysis of tissues of a healthy donor for IgD, AIRE, CD19 and DNA. Arrow heads indicate follicular IgD^+^ plasmablasts. Dotted lines mark the boundary between follicular mantle zone and the follicle. Bars: 40 μm (C) and 15 μm (D). (E) Flow cytometric analysis of AIRE expression in tonsillar total CD19^+^ B cells, IgD^+^CD38^−^naive B cells, IgD^+^CD38^+^ founder GC (FGC) B cells, IgD^−^CD38^+^ GC B cells and IgD^−^CD38^−^memory B cells. The data represent 5 donors. (F and G) Flow cytometric and statistical analyses of Aire (GFP) expression in splenic and ILN viable CD19^+^B220^+^FAS^+^GL7^+^ GC B cells, CD19^+^B220^+^FAS^−^GL7^−^ non-GC B cells and CD19^lo^B220^lo^CD138^+^ PCs of B6 mice (shaded histograms, *n* = 5 for spleen and *n* = 4 for ILN) or B6.*Aire^Adig^* mice (colored histograms, *n* = 5 for spleen and *n* = 4 for ILN) after 4 i.p. immunizations with NP_32_-KLH with CFA and subsequently with IFA. Data are represented as mean ± SEM. \*\**P* < 0.01, \*\*\**P* < 0.001, by 1-way ANOVA with Tukey’s post hoc test. (H) qPCR analysis of *Aire* transcript levels in CD19^+^B220^+^FAS^−^GL7^−^ non-GC B cells (*n* = 5) and CD19^+^B220^+^FAS^+^GL7^+^GFP^+^ Aire-expressing GC B cells (*n* = 4) of *Aire^Adig^* mice after 1 i.p. immunization with SRBC and CFA. Data are represented as mean ± SEM. \*\*\**P* < 0.001, by 2-tailed unpaired *t*-test. (I) Flow cytometric and statistical analyses of CD83^+^CXCR4^lo^ LZ and CD83^−^CXCR4^hi^ DZ B cells in splenic total GC and GFP^+^ GC B cells of immunized B6.*Aire^Adig^* mice, by 2-tailed paired *t*-test. (J and K) Immunofluorescence analysis of the tonsillar tissue of a healthy donor for IgD, CD23, Pax5 and AIRE. The dotted line outlines the follicles and delineates the border between LZ and DZ. Bars: 200 μm (J) or 30 μm (K).

Similar to human B cells, AIRE was found in B cells in the splenic follicles of immunized mice (**Figure S1E** and **F**). Consistently, in the *Aire^Adig^*reporter mice (Gardner et al., 2008) after intraperitoneal immunizations, B cell AIRE expression was detected in FAS^+^GL7^+^ GC B cells in the spleen, inguinal lymph nodes (ILNs), mesenteric lymph nodes (MLNs) and Peyer’s patches (PPs) and in thymic B cells, but not in FAS^−^GL7^−^ non-GC B cells or CD138^+^ PCs in these tissues, or in peripheral blood or peritoneal B cells (**Figure 1F** and **G**, **Figure S1G–J**). Consistently, *Aire* transcript level was markedly higher in splenic GFP^+^ GC B cells than non-GC B cells (**Figure 1H**). Furthermore, AIRE was detected in both CXCR4^+^CD83^−^ dark zone (DZ) and CXCR4^lo^CD83^+^ light zone (LZ) B cells in mouse spleens (**Figure 1I**) and DZ and LZ B cells in human tonsils (**Figure 1J** and **K**). Altogether, these data indicate that AIRE expression in GC B cells is a specific and conserved characteristic of human and mouse secondary lymphoid organs.

### Follicular B Cell AIRE Expression Requires CD40 Signaling

The induction of T cell-dependent GC B cell responses in secondary lymphoid organs involves B cell antigen presentation to primed cognate T helper (T_H_) cells and clonal proliferation upon receiving T_H_ cell signals, such as CD40-ligand (CD40L) and interleukin (IL)-4 (Liu et al., 1989; Yusuf et al., 2010). CD40 signaling was previously reported to promote AIRE expression by mTECs and thymic B cells (Akiyama et al., 2008; Yamano et al., 2015). To determine the regulation of AIRE expression in follicular B cells, we examined the tonsillar tissue of a patient with the rare primary immunodeficiency (PID) Hyper-IgM Syndrome type 3 (HIGM3), which is caused by loss-of-function mutations in the *CD40* gene (Durandy et al., 2005). In contrast to the prominent AIRE expression in tonsillar follicular B cells of healthy subjects (**Figure 1A** and **B**, **Figure S1B** and **C**), the tonsil of the HIGM3 patient harbored enlarged follicles containing B cells that had no expression of AIRE and failed to downregulate cell surface IgD (**Figure 2A** and **B**). Of note, tonsillar IgD plasmablasts were absent from the follicles but still present in the extrafollicular area, which is consistent with the existence of T cell-independent pathways for their generation as we reported previously (Chen et al., 2009), and indicates a specific shutdown of the T cell-dependent GC antibody diversification machinery. These results demonstrate a requirement for CD40 signaling in follicular B cell AIRE expression *in vivo*.

**Figure 2.**
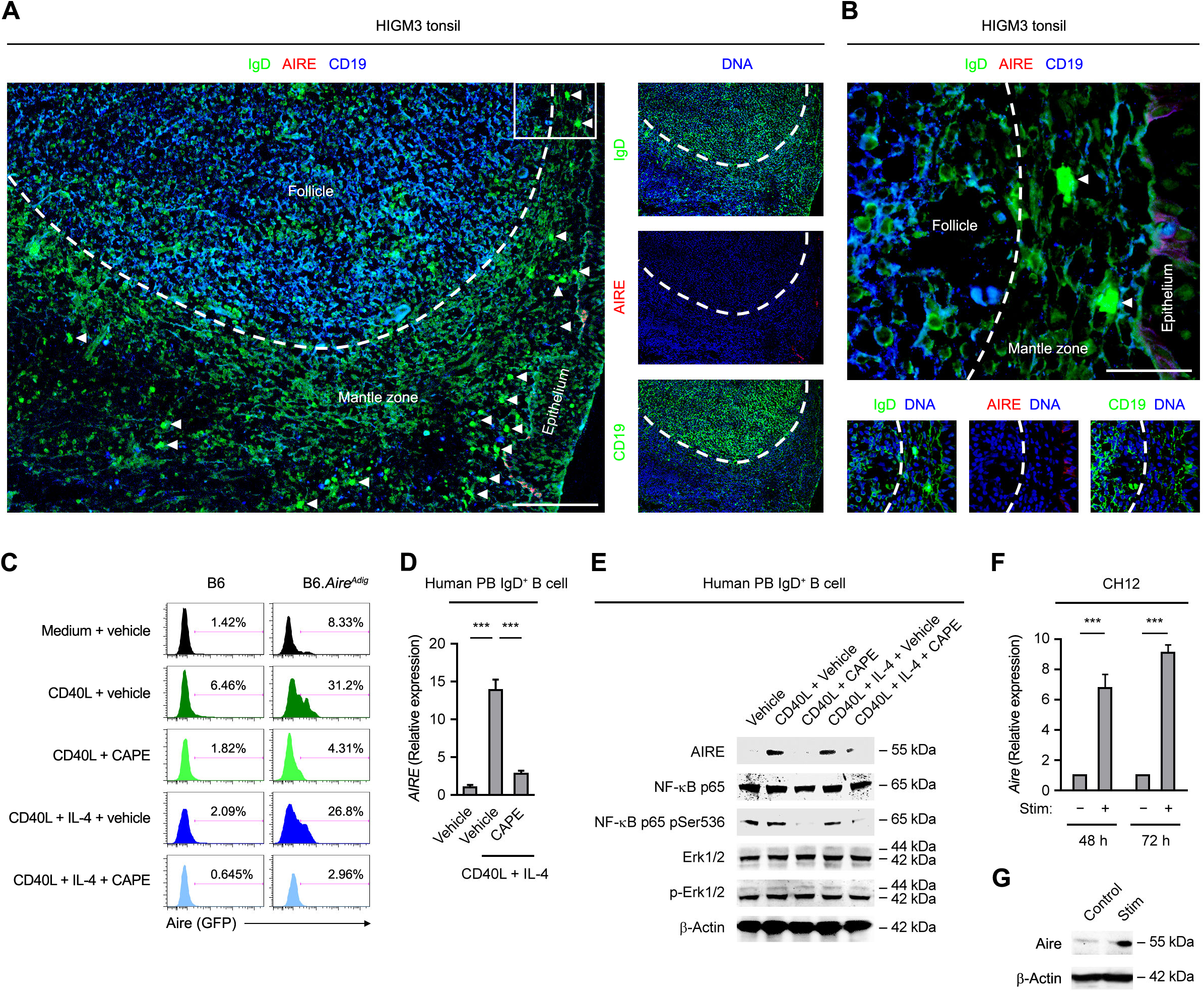
Follicular B Cell AIRE Expression Requires CD40 Signaling. (A and B) Immunofluorescence analysis of tonsillar tissues of a HIGM3 patient for IgD, AIRE, CD19 and DNA. The area outlined in A is shown with a higher magnification in B. The dotted line outlines the follicles. Bars: 100 μm (A) or 25 μm (B). (C) Flow cytometric analysis of Aire (GFP) expression in splenic B cells of a B6 or B6.*Aire^Adig^* mouse treated for 3 d with medium or CD40L with or without IL-4 in the absence (vehicle) or presence of CAPE. The data represent the results from 3 B6 and 3 B6.*Aire^Adig^* mice. (D and E) qRT-PCR and Western Blot analyses of *AIRE* transcript and protein levels, the protein levels of total and Ser536-phosphorylated NF-κB p65, as well as total and Thr202/Tyr204-phosphorylated Erk1/2 in human peripheral blood IgD^+^ B cells treated with medium or CD40L, or CD40L and IL-4, in the presence of vehicle or CAPE for 3 d. Data are represented as mean ± SEM. \*\*\**P* < 0.001, by 2-tailed unpaired *t*-test. (F and G) qRT-PCR and Western Blot analyses of *Aire* transcript and Aire protein levels in mouse CH12 cells treated with anti-CD40, TGF-β and 100 ng/ml IL-4 for 3 d. Data are represented as mean ± SEM. *P* < 0.001, by 2-tailed unpaired *t*-test.

To further ascertain a role of CD40 signaling in promoting B cell AIRE expression, we treated mouse splenic naive resting B cells and human peripheral blood IgD^+^ mature naive B cells *in vitro* with CD40L alone or in combination with IL-4. Consistent with the above *in vivo* data, these B cells expressed AIRE protein and transcript upon CD40L stimulation (**Figure 2C–E**). AIRE induction was abrogated by caffeic acid phenethyl ester (CAPE), a selective inhibitor of nuclear factor-kappa B (NF-κB) (Natarajan et al., 1996), a transcription factor activated by CD40 (**Figure 2C–E**). Furthermore, AIRE transcript and protein were also induced in the mouse B cell line CH12 (Nakamura et al., 1996) upon anti-CD40 stimulation (**Figure 2F** and **G**). Collectively, these results show that CD40 signaling promotes AIRE expression in GC B cells *in vivo* and in B cells and cell lines *in vitro*.

### AIRE in B Cells Inhibits CSR and SHM

B cell activation by antigens results in antibody diversification such as CSR, which primarily occurs prior to GC entry (Roco et al., 2019), and SHM in the GC (Murphy and Weaver, 2016b), we sought to determine the B cell-intrinsic function of AIRE in CSR and SHM. We employed several independent experimental approaches to achieve this goal.

Firstly, we engrafted lethally irradiated B cell-deficient μMT mice with the BM of CD45.1^+^ *Aire*^+/+^ and CD45.2^+^ *Aire*^−/−^ mice depleted of B220^+^ cells at a ratio of 1:1 (**Figure S2A**), allowing the donor B cell compartment to develop in the same environment in which the thymic epithelium is *Aire*^+/+^ so that the autoimmune manifestations of AIRE deficiency do not develop. Twenty-eight days later, naive resting splenic unswitched *Aire*^+/+^ and *Aire*^−/−^ B cells from these BM chimeras were adoptively transferred to secondary μMT recipients at a ratio of 1:1 (**Figure S2B**). These secondary μMT recipients were subsequently immunized with the T cell-dependent antigen (4-hydroxy-3-nitrophenylacetyl)_32_-Keyhole Limpet Hemocyanin (NP_32_-KLH) and analyzed the affinity maturation and class switching of NP-specific donor B cells by flow cytometry (**Figure 3A**). After the immunizations, the splenic GC B cell compartment contained more *Aire*^−/−^B cells than *Aire*^+/+^ B cells (**Figure 3B**), and the GC *Aire*^−/−^ B cells had a higher NP_8_-to-NP_36_ binding ratio and a higher percentage of NP_8_-binding cells in total NP-specific cells than their *Aire*^+/+^ counterpart (**Figure 3C** and **D**), indicating increased affinity maturation. These data suggest that AIRE inhibits SHM in a B cell-intrinsic manner.

**Figure 3.**
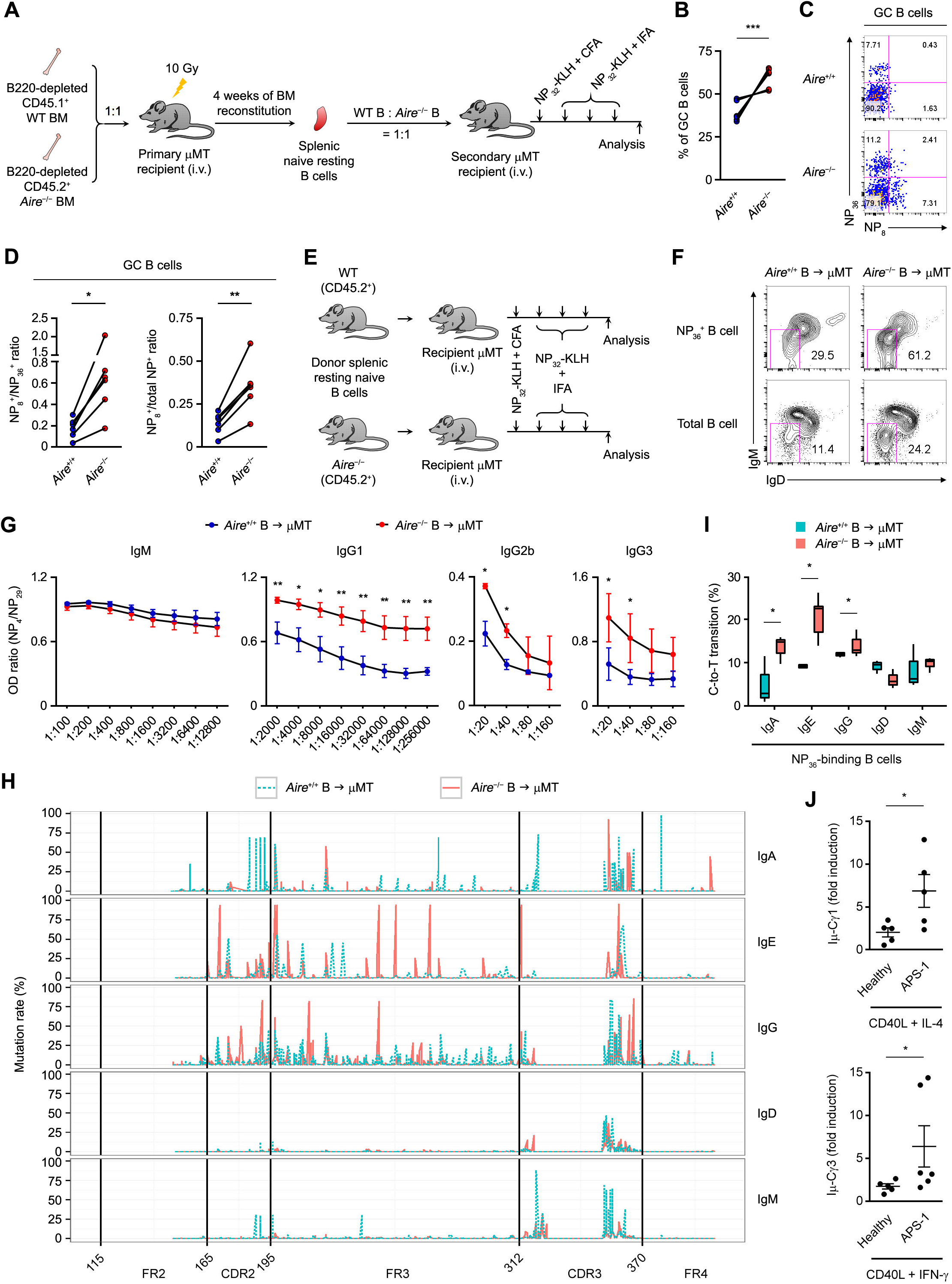
AIRE Inhibits CSR and SHM in B cells. (A) The generation and immunization of *Aire*^+/+^ and *Aire*^−/−^ BM and B cell chimeric mice. (B) The percentage of CD45.1^+^ *Aire*^+/+^ and CD45.2^+^ *Aire*^−/−^ B cells in the splenic GC B cells of the secondary μMT recipient mice (*n* = 6) after the immunizations. \*\*\**P* < 0.001, by 2-tailed paired *t*-test. (C and D) Flow cytometric and statistical analyses of the ratio of NP_8_- to NP_36_-binding (C) and NP_8_- to total NP-binding (D) GC B cells after immunizations of the secondary μMT recipient mice (*n* = 6). \*\**P* < 0.05, \*\**P* < 0.01, by 2-tailed paired *t*-test. (E) The generation and immunization of *Aire*^+/+^ and *Aire*^−/−^ B cell chimeric mice. (F) Flow cytometric analysis of surface IgD and IgM on NP_36_-binding B cells in μMT recipients of *Aire*^+/+^ or *Aire*^−/−^ B cells immunized with NP_32_-KLH. The result represents 3 age- and sex-matched μMT recipients each of B cells from 3–5 age- and sex-matched littermate donor *Aire*^+/+^ or *Aire*^−/−^ mice. (G) The ratios of the titers of circulating NP_4_-binding to NP_29_-binding IgM, IgG1, IgG2b and IgG3 in immunized μMT recipient mice of *Aire*^+/+^ or *Aire*^−/−^ B cells. The results represent 4 experiments, each consisting of B cells from 3–5 age- and sex-matched littermate donor mice and 6–8 age- and sex-matched littermate μMT recipient mice. \**P* < 0.05, \*\**P* < 0.01, by 2-tailed unpaired *t*-test. (H) The SHM landscape across IgHV, including FR2, CDR2, FR3, CDR3 and FR4, of NP_36_-binding IgM^‒^IgD^‒^ or IgM^+^IgD^+^ *Aire*^+/+^ and *Aire*^‒/‒^ donor B cells in μMT recipients after immunizations with NP_32_-KLH. The result represents 3 μMT recipients of *Aire*^+/+^ donor B cells and 3 μMT recipients of *Aire*^‒/‒^ donor B cells. (I) Frequencies of C-to-T transitions in SHMs in IgHV of NP-specific IgG^+^, IgA^+^ or IgE^+^ splenic B cells from μMT recipient mice of *Aire*^+/+^ or *Aire*^‒/‒^ B cells after immunizations with NP_32_-KLH. Data are represented as median ± upper/lower quartile. \**P* < 0.05, \*\**P* < 0.01, by 1-tailed unpaired *t*-test. (J) qRT-PCR analysis of the fold induction of Iì-Cã1 and Iì-Cã3 post-switch transcript levels in peripheral blood IgD^+^CD27^‒^ naïve B cells from healthy subjects (*n* = 5) or APS-1 patients (*n* = 5) stimulated for 3 d with CD40L and IL-4 or IFN-γ over the respective unstimulated control B cells. \**P* < 0.05, by 1-tailed unpaired *t*-test (upper panel) or 1-tailed Mann-Whitney *U* test (lower panel).

Secondly, we adoptively transferred an equal number of naive resting unswitched B cells from CD45.2^+^ *Aire*^+/+^ or CD45.2^+^ *Aire*^−/−^ donor mice into μMT recipients and assessed the CSR and SHM of NP-specific donor B cells after the immunizations by flow cytometry, enzyme-linked immunosorbent assay (ELISA) and Ig heavy (IgH) chain variable (V) region mutation profiling with next-generation sequencing (**Figure 3E**). The *Aire*^+/+^ and *Aire*^−/−^ donor B cells exhibited a similar phenotype and had a comparable NP_8_-to-NP_36_ binding ratio before the transfer (**Figure S2C–E**). Following the immunizations, *Aire*^+/+^ and *Aire*^−/−^ donor B cells entered the GC similarly in their respective recipients (**Figure S2F**), and GC *Aire*^+/+^ and *Aire*^−/−^ donor B cells exhibited similar expression of major co-stimulatory and co-inhibitory molecules (**Figure S2G**). µMT recipients of *Aire*^+/+^ or *Aire*^−/−^ B cells had a similar proportion of CXCR5^+^PD-1^+^ follicular helper T (T_FH_) cells (**Figure S2H**) and Foxp3^+^CD25^+^ follicular regulatory T (T_FR_) cells in the spleen (**Figure S2I**). However, NP-specific *Aire*^−/−^ donor B cells exhibited elevated class switching by harboring a much higher fraction of IgM^−^IgD^−^ cells than NP-specific *Aire*^+/+^ donor B cells (**Figure 3F**), and underwent increased affinity maturation by producing IgG1, IgG2b and IgG3 of higher NP_4_-to-NP_29_ binding ratios (**Figure 3G**). In contrast, a higher NP_4_-to-NP_29_ binding ratio was not observed in the IgM compartment (**Figure 3G**). Increased CSR and SHM of donor *Aire*^−/−^ B cells were not a secondary effect of differential AIRE expression by donor B cells in the thymus, because we found a negligible number of B cells in the thymus of the μMT recipient mice and the adoptive transfer did not lead to an increase in B cell numbers in the thymic stroma of the recipients at the end of the experiment, indicating that the donor B cells did not enter the thymus of the recipients during the course of our experiment (**Figure S2J** and **K**). Analysis of the IgH variable region (IgHV) SHM landscape of NP-specific *Aire*^+/+^ and *Aire*^−/−^ donor B cells sorted from the recipient mice (**Figure S2L**) showed that *Aire*^−/−^ NP-specific class-switched IgG and IgE B cells had higher rates of IgHV SHMs in complementarity-determining region 2 (CDR2) and framework region 3 (FR3) than *Aire*^+/+^ NP-specific B cells (**Figure 3H**). Importantly, such an increased SHM profile was not seen in the NP-specific IgM and IgD compartments (**Figure 3H**). In addition, there was an increased frequency of C-to-T transitions among the SHMs in the IgH V region coding sequences in *Aire*^−/−^ NP-specific B cells compared to *Aire*^+/+^ NP-specific B cells (**Figure 3I**), which is a molecular signature associated with the action of AID in the IgH V region (Maul et al., 2011). *Aire*^+/+^ and *Aire*^−/−^splenic B cells exhibited comparable proliferation and apoptosis upon stimulation *ex vivo* (**Figure S3A–D**). These data suggest that AIRE suppresses CSR and SHM in AID-experienced B cells.

Thirdly, we sorted IgD^+^CD27^−^ mature naive B cells from the peripheral blood of healthy subjects and APS-1 patients and compared their CSR *in vitro* upon stimulation with T cell-dependent stimuli. Naive B cells of APS-1 patients underwent increased CSR compared to those of healthy subjects by expressing higher levels of the post-switch transcript Iμ-Cγ1 or Iμ-Cγ3 following stimulation with CD40L and IL-4 or CD40L and interferon (IFN)-γ, respectively (**Figure 3J**).

Finally, using CRISPR-*Cas9*-mediated gene editing, we disrupted the *Aire* gene in the murine B cell line CH12, which can undergo class switching from IgM to IgA upon stimulation with anti-CD40, TGF-β and IL-4 (Nakamura et al., 1996). We identified 3 *Aire*^−/−^ CH12 clones with frame-shift mutations in both *Aire* alleles (**Table S1**, and **Figure S4A** and **B**) that were devoid of AIRE protein expression (**Figure S4C**). Upon stimulation, these *Aire*^−/−^ CH12 clones underwent elevated IgA CSR (**Figure 4A** and **B**) with concomitantly increased levels of the Iα-Cµ circle transcript compared to their parental *Aire*^+/+^ CH12 cells (**Figure 4C**). The increased CSR was specific to IgA, as Iγ1-Cµ, the circle transcript of IgG1, an isotype that CH12 cells do not switch to, was not affected by *Aire* deficiency (**Figure 4D**). Exaggerated IgA CSR in *Aire*^−/−^ CH12 cells was not a result of increased induction of AID (**Figure 4E** and **F**) or germline transcription (**Figure 4G**), nor a result of increased survival (**Figure S4D** and **E**). Remarkably, WT AIRE suppressed cytokine-induced CSR when re-introduced into *Aire*^−/−^ CH12 cells (**Figure 4H**). Collectively, these results demonstrate that B-cell intrinsic AIRE negatively regulates CSR and SHM.

**Figure 4.**
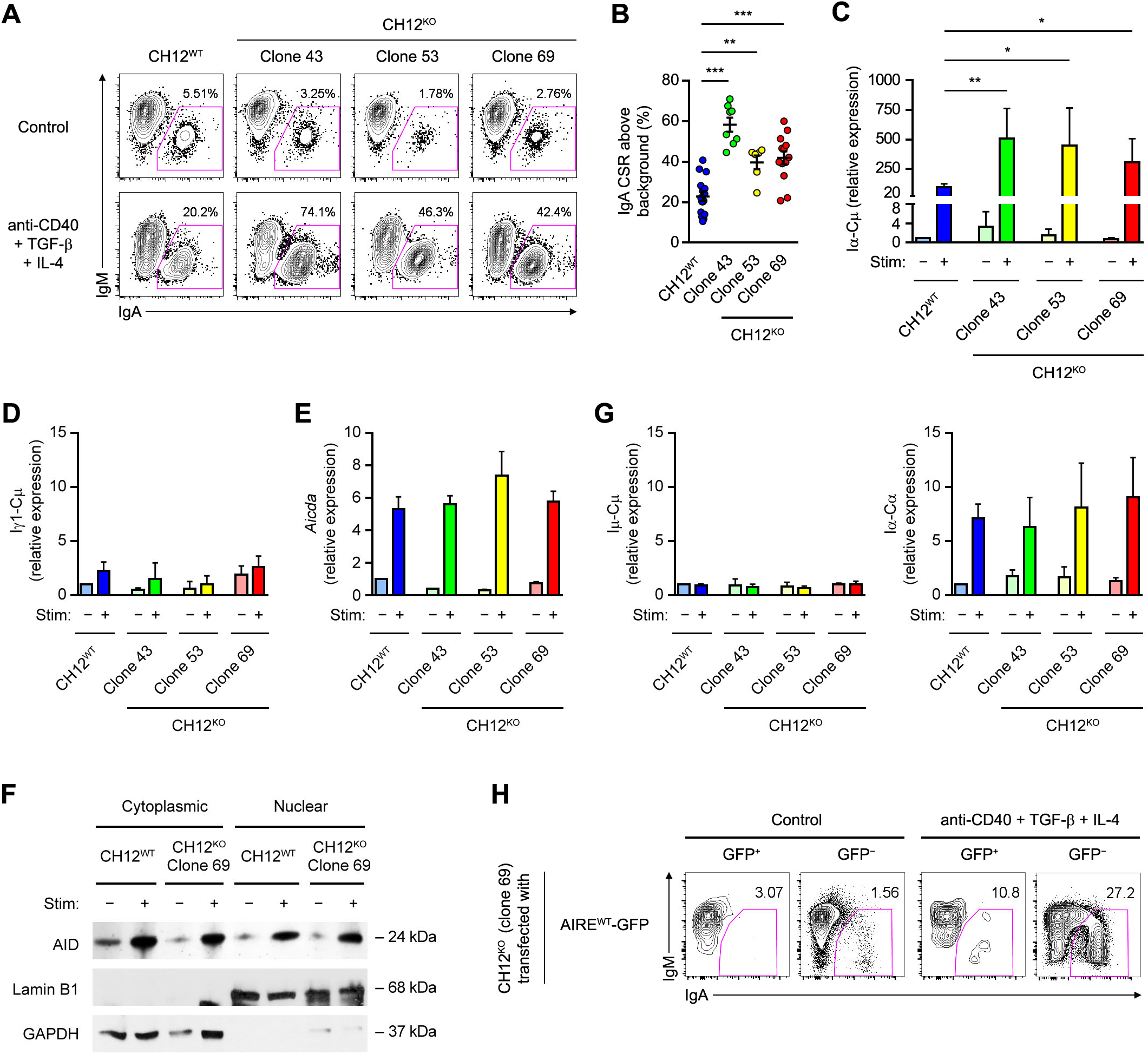
*Aire*^−/−^ CH12 Cells Undergo Elevated CSR. (A and B) Flow cytometric and statistical analyses of IgA CSR in WT and *Aire*^−/−^ CH12 cells treated with medium (Control) or anti-CD40, TGF-β and IL-4 for 3 d. Data in **F** was determined as the difference between the percentages of IgA^+^IgM^−^ cells in stimulated samples to unstimulated samples. The results (mean ± SEM) represent or compare 16 experiments involving WT CH12 cells, 8 experiments involving clones 43, 6 experiments involving clone 53, and 13 experiments involving clone 69. \*\**P* < 0.01, \*\*\**P* < 0.001, by 2-tailed unpaired *t*-test. (C) qRT-PCR analysis of the Iα-Cμ circle transcript levels (mean ± SEM) in WT and *Aire*^−/−^ CH12 cells treated with medium (Control) or anti-CD40, TGF-β and IL-4 for 3 d. The results compare 3 experiments. \**P* < 0.05, \*\**P* < 0.01, by 1-tailed unpaired *t*-test. (D) qRT-PCR analysis of the Iγ1-Cμ circle transcript level in *Aire*^+/+^ CH12 cells and *Aire*^−/−^ CH12 cell clones 43, 53 and 69 that were either unstimulated or stimulated with anti-CD40, TGF-β and IL-4 for 3 d. The result was normalised using the respective *Actb* transcript level and expressed as fold of induction (mean ± SEM) relative to unstimulated *Aire*^+/+^ CH12 cells. (E and F) Western Blot analysis of AID in WT and *Aire*^‒/‒^ CH12 cells that were either unstimulated or stimulated with anti-CD40, TGF-β and IL-4 for 3 d. Lamin B1 and GAPDH were used as the control for nuclear and cytoplasmic proteins, respectively. The data are presented as mean ± SEM and represent 3 experiments. (G) qRT-PCR analysis of *Aicda* and the Iμ-Cμ and Iα-Cα germline transcript levels (mean ± SEM) in *Aire*^+/+^ CH12 cells and *Aire*^−/−^ CH12 cell clones 43, 53 and 69 that were either unstimulated or stimulated with anti-CD40, TGF-β and IL-4 for 3 d. (H) Flow cytometric analysis of IgA CSR in *Aire*^−/−^ CH12 cells (clone 69) transfected with a plasmid expressing WT (AIRE^WT^-GFP) AIRE-GFP and treated with medium (Control) or anti-CD40, TGF-β and IL-4 for 3 d. The results represent 3 experiments.

### AIRE Interacts with AID in GC B Cells

We next investigated the mechanism by which AIRE inhibits peripheral antibody diversification. Given AID as the obligatory enzyme that mediates CSR and SHM (Muramatsu et al., 2000), we asked whether AIRE inhibits AID function in B cells. To this end, we first examined whether AIRE and AID interact in GC B cells. AIRE and AID co-localized in the nuclei of tonsillar IgD^−^CD38^+^ GC B cells (**Figure 5A** and **B**, and **Figure S5A**) but not in IgD^+^CD38^−^ naive B cells (**Figure S5B**), IgD^−^CD38^−^ switched memory B cells (**Figure S5C**) or IgD^−^CD38^hi^ switched PCs (**Figure S5D**), albeit a low level of nuclear AIRE and AID were detected in a small fraction of IgD^+^CD38^+^ FGC B cells (**Figure S5E**). Using an AID antibody validated for immunoprecipitation (IP) and Chromatin IP (ChIP) (Vuong et al., 2009), we found that AIRE interacted with AID in human tonsillar CD19^+^ and CD19^+^IgD^−^ cell fractions (**Figure 5C**). AID and AIRE did not interact via DNA in GC B cells as they still co-immunoprecipitated after DNAse I treatment (**Figure S6A**). AIRE also co-immunoprecipitated with AID in splenic B cells of immunized WT but not *Aire*^−/−^or *Aicda*^−/−^ mice (**Figure 5D** and **Figure S6B**). Based on these data, we conclude that AIRE interacts with AID in GC B cells *in vivo*.

**Figure 5.**
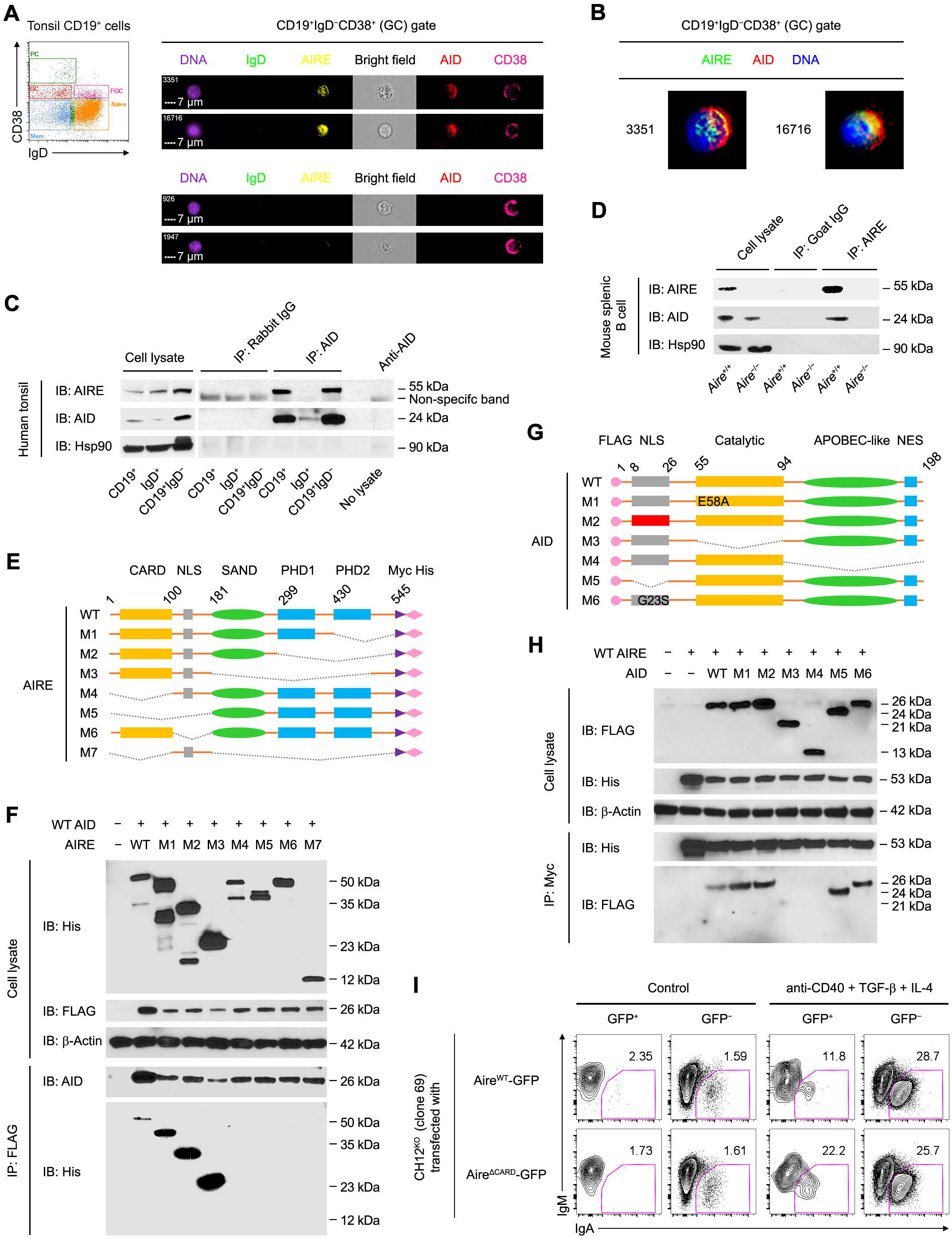
AIRE Interacts with AID in GC B Cells. (A and B) Imaging flow cytometric analysis of AIRE and AID in tonsillar IgD^−^CD38^+^ GC B cells of a healthy donor. Bars: 7 μm (A). The results represent 3 donors. (C and D) Co-IP of AIRE and AID in tonsillar CD19^+^ total, IgD^+^ naive and FGC and CD19^+^IgD^‒^GC and memory B cells of a healthy donor, and in splenic CD19^+^ B cells of a B6 mouse after 3 doses of immunization with SRBCs. The results are representative of tonsils of 4 donors and spleens of 3 mice. (E) The domain structures of recombinant WT and mutant human AIRE and AID molecules. Dotted lines indicated the deleted regions in the mutant proteins. (F) Co-IP of WT AID and WT or mutant AIRE in HKB-11 cells 24 h after transfection of plasmid(s) encoding WT AID and WT or mutant AIRE proteins. (G) The domain structures of recombinant WT and mutant human AID proteins. (H) Co-IP of WT AIRE and WT or mutant AID in HKB-11 cells 24 h after transfection of plasmid(s) encoding WT AIRE and WT or mutant AID proteins. The results in E and G are representative of 3 experiments. (I) Flow cytometric analysis of IgA CSR in *Aire*^−/−^ CH12 cells (clone 69) transfected with a plasmid expressing either WT (Aire^WT^-GFP) or CARD-deficient (Aire^ΔCARD^-GFP) AIRE-GFP and treated with medium (Control) or anti-CD40, TGF-β and IL-4 for 3 d. The results represent 3 experiments.

### Interaction of AIRE with AID Requires the CARD and NLS domains of AIRE

We subsequently generated a series of deletion mutants of AIRE with C-terminal Myc and His tags to characterize AIRE’s interaction with AID (**Figure 5E** and **Table S2A**). AIRE mutants missing the N-terminal caspase activation and recruitment domain (CARD) and/or nuclear localization signal (NLS) lost the ability to interact with AID (**Figure 5F**), suggesting the requirement for the CARD domain and nuclear localization of AIRE for interaction with AID. Furthermore, using a series of deletion, domain replacement or point mutants of AID with an N-terminal FLAG tag (**Figure 5G** and **Table S2B**), we found that the interaction between AID and AIRE required both the catalytic and APOBEC-like domains of AID, although the catalytic activity of AID was not necessary, as the catalytically inactive AID^E58A^ mutant (Patenaude et al., 2009) still interacted with AIRE (**Figure 5H**). The AID point mutation G23S, which substantially abrogates the SHM but not much CSR activity of AID (Wei et al., 2011), did not affect the interaction with AIRE (**Figure 5H**). Echoing the CARD-dependent interaction of AIRE with AID, a CARD domain deletion mutant of AIRE had impaired ability to suppress CSR when introduced into *Aire*^−/−^ CH12 cells (**Figure 5I**).

### AIRE Inhibits the Function of AID during Antibody Diversification

Since AID enzymatically generates uracils in DNA (Bransteitter et al., 2003; Chaudhuri et al., 2003; Sohail et al., 2003), we employed a genomic uracil dot blot assay to directly test the effect of AIRE on the activity of AID (**Figure 6A** and **B**). In CH12 cells, increased uracils were detected in the genome upon stimulation with anti-CD40, TGF-β and IL-4 to undergo IgA CSR, which peaked on day 3 (Shalhout et al., 2014), whereas *Acida*^−/−^ CH12 cells failed to accumulate genomic uracils after stimulation (**Figure 6C**), indicating that the emergence of genomic uracils was mediated by AID. Upon stimulation, *Aire*^−/−^ CH12 cells harbored higher levels of genomic uracil than *Aire*^+/+^ CH12 cells (**Figure 6C** and **D**). This result reflects the inhibition of AID’s enzymatic function by AIRE at a step upstream of the deamination reaction.

**Figure 6.**
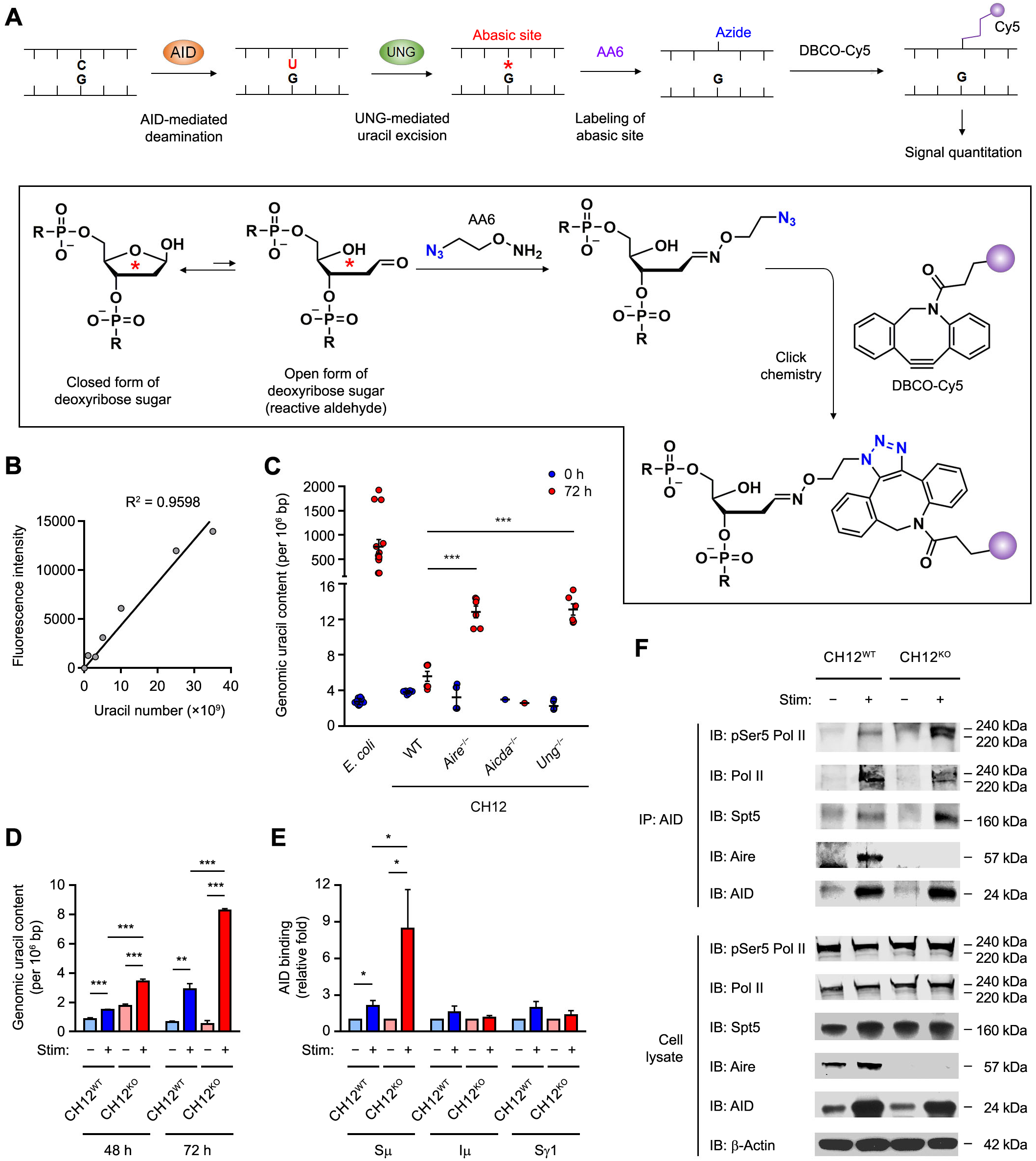
AIRE Inhibits AID Function by Interfering with AID Targeting to Its IgH DNA Substrate. (A and B) The principle, chemistry and calibration of the dot blot assay for the quantitation of genomic uracil content. (C) The genomic uracil levels in WT, *Aire*^‒/‒^, *Aicda*^‒/‒^, or *Ung*^‒/‒^ CH12 cells after 72 h of treatment without or with anti-CD40, TGF-β and IL-4. The results are presented as mean ± SEM and represent 3 experiments. \*\*\**P* < 0.001, by 2-tailed unpaired *t*-test. Bisulfite-treated *E. coli* DNA was included as a positive control. (D) The genomic uracil content in WT and *Aire*^‒/‒^ CH12 cells after 48 or 72 h of treatment without or with anti-CD40, TGF-β and IL-4. The results are presented as mean ± SEM and represent 3 experiments. \*\**P* < 0.01, \*\*\**P* < 0.001, by 2-tailed unpaired *t*-test. (E) ChIP-qPCR analysis for the interaction of AID with Sμ, Iμ and Sγ1 regions in WT and *Aire*^‒/‒^CH12 cells after 72 h of treatment without or with anti-CD40, TGF-β and IL-4. The results are presented as mean ± SEM and represent 3 experiments. \**P* < 0.05, by 2-tailed unpaired *t*-test. (F) Co-IP of AID with pSer5-Pol II, total Pol II, Spt5 and Aire in WT and *Aire*^‒/‒^ CH12 cells after 72 h of treatment without or with anti-CD40, TGF-β and IL-4. The results represent 3 experiments.

We further investigated how AIRE may inhibit AID’s enzymatic activity. Upstream of DNA deamination by AID is the proper targeting of AID to the IgH S regions at sites of Pol II stalling (Chaudhuri et al., 2003; Pavri et al., 2010). We hypothesized that AIRE may interfere with the targeting of AID to IgH S regions during CSR. Using ChIP and IP assays, we found increased AID binding to the Sµ, but not Iµ or Sγ1, region (**Figure 6E**) and increased interaction of AID with serine-5 phosphorylated Pol II and its associated factor Spt5 at a promoter-proximal pause site (Peterlin and Price, 2006) in stimulated *Aire*^−/−^ CH12 cells compared to stimulated *Aire*^+/+^ CH12 cells (**Figure 6F** and **Figure S6C**). These data suggest that AIRE inhibits AID function in B cells by interfering with the interaction of AID with transcriptionally paused Pol II and the targeting of AID to its IgH DNA substrate.

### AIRE Deficiency in B Cells Engenders Humoral Autoimmunity and Compromises Skin *Candida* Immune Defense

Given that the vast majority of APS-1 patients mysteriously develop chronic mucocutaneous candidiasis (CMC) as an early clinical symptom (Kisand and Peterson, 2015), we sought to determine the functional impact of B cell-intrinsic AIRE in humoral immunity and anti-*Candida* immune defense, and asked whether AIRE deficiency in peripheral B cells could promote APS-1-like CMC. We employed a mouse dermal candidiasis model which allows skin *C. albicans* infection to be established without the use of immunosuppressive agents (Conti et al., 2014). μMT recipient mice reconstituted with either *Aire*^+/+^ or *Aire*^−/−^ B cells were first exposed to heat-killed *C. albicans* pseudohyphae and subsequently infected cutaneously with live *C. albicans* pseudohyphae (**Figure 7A**). Four days after infection, μMT recipient mice of *Aire*^−/−^ B cells had heightened fungal burden in the skin (**Figure 7B** and **C**) and concomitant elevation of circulating autoantibodies to IL-17A, IL-17F and IL-22 as compared to μMT recipients of *Aire*^+/+^ B cells (**Figure 7D** and **Figure S7A**). The sera of μMT recipients of *Aire*^−/−^ B cells contained enhanced blocking activity of the binding of IL-17A, IL-17F and IL-22 to their receptors (**Figure 7E**). In addition, the dermal infection site of μMT recipients of *Aire*^−/−^ B cells harbored reduced IL-17A-and IL-22-producing T cells (**Figure 7F** and **G**), which was accompanied by diminished neutrophils infiltration into the infected skin tissue (**Figure 7H**). These results are reminiscent of the aberrant production of class-switched neutralizing autoantibodies against T_H_17 cytokines in APS-1 patients that may impair anti-*C. albicans* immunity (Kisand et al., 2010; Meyer et al., 2016a; Puel et al., 2010), and collectively show that AIRE deficiency in peripheral B cells compromises cutaneous anti-*Candida* immune defense and promotes APS-1-like CMC by engendering humoral autoimmunity.

**Figure 7.**
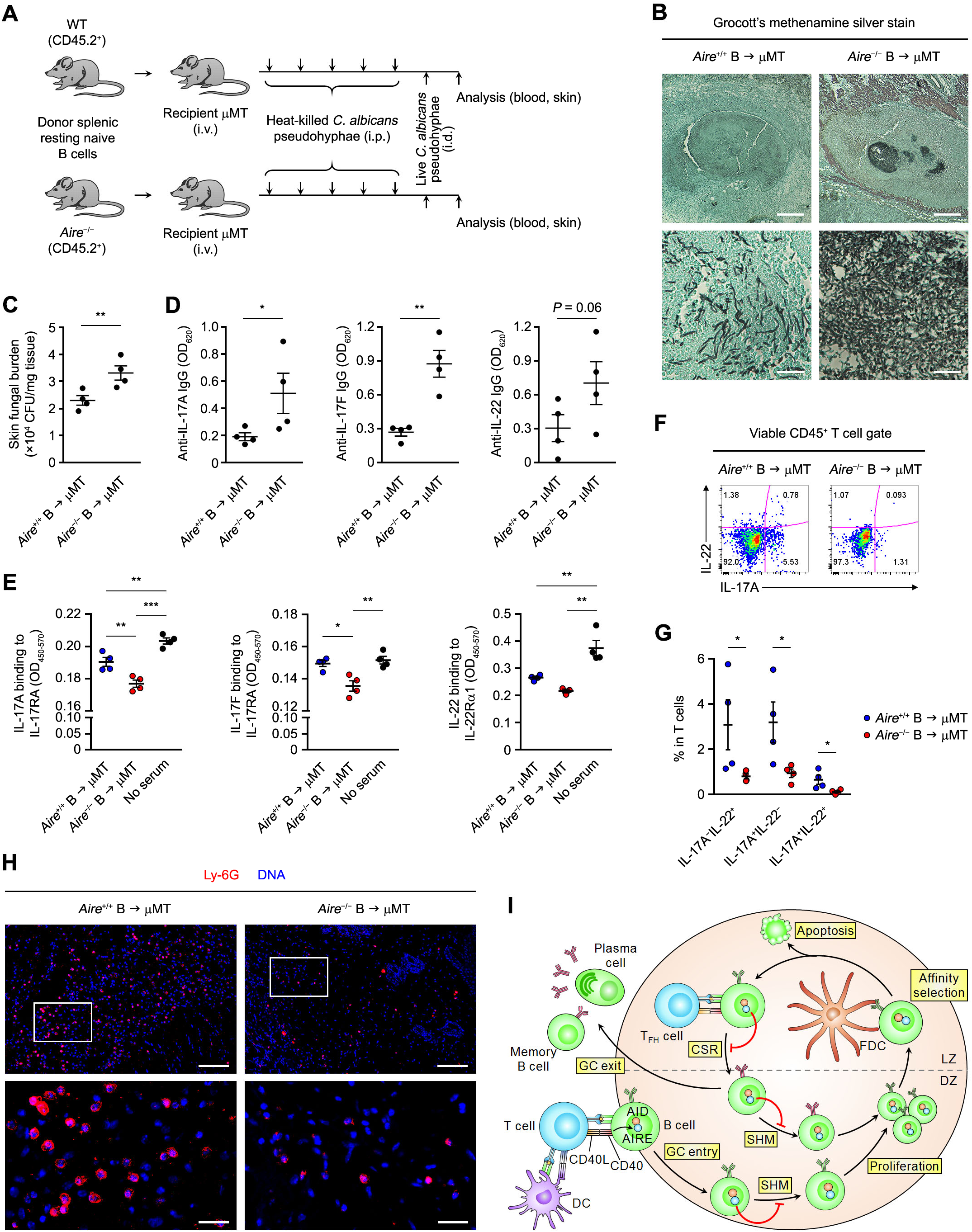
Aire Deficiency in B Cells Promotes Humoral Autoimmunity and Compromises Cutaneous Anti-*Candida* Defense. (A) The cutaneous candidiasis infection model. (B and C) GMS stain of cutaneous *C. albicans* and skin fungal burden (CFU per mg of tissue) (mean ± SEM) in μMT recipient mice of *Aire*^+/+^ or *Aire*^‒/‒^ donor B cells 4 d after infection. Bars: 1 mm (B, upper panels) or 100 μm (B, lower panels). \*\**P* < 0.01, by 1-tailed unpaired *t*-test. (D) ELISA of the levels (mean ± SEM) of autoantibodies binding to IL-17A, IL-17F and IL-22 in the sera of μMT recipient mice of *Aire*^+/+^ or *Aire*^‒/‒^ donor B cells 4 d after infection. \**P* < 0.05, \*\**P* < 0.01, by 1-tailed unpaired *t*-test. (E) ELISA of the levels (mean ± SEM) of blocking activity of IL-17A, IL-17F and IL-22 to their receptors in the sera of μMT recipient mice of *Aire*^+/+^ or *Aire*^‒/‒^ donor B cells 4 d after infection. \**P* < 0.05, \*\**P* < 0.01, \*\*\**P* < 0.001, by 1-way ANOVA with Tukey’s post hoc test. (F and G) Flow cytometric and statistical analyses of IL-17A and IL-22 expression in cutaneous T cells of μMT recipient mice of *Aire*^+/+^ or *Aire*^‒/‒^ donor B cells (*n* = 4 in each group) 4 d after infection and after *ex vivo* re-stimulation. The data are represented as mean ± SEM. \**P* < 0.05, by 1-tailed unpaired *t*-test. (H) Immunofluorescence analysis of Ly-6G (red) and DNA (blue) in cutaneous tissues surrounding the *C. albicans* infection site in μMT recipient mice of *Aire*^+/+^ or *Aire*^‒/‒^ donor B cells 4 d after infection. The results in B–H represent 2 experiments, with 4 mice per group in each experiment. Bars: 160 μm (upper panels) or 40 μm (lower panels). (I) A simplified schematic of AIRE-mediated GC checkpoint of antibody diversification in B cells. At the T–B cell border of secondary lymphoid organs, B cells present antigens to and receive co-stimulation from DC-activated T cells, which also induce AIRE expression in B cells via CD40. The activated B cells enter the GC DZ and undergo SHM, proliferation and subsequent affinity selection by interacting with antigens on the surface of follicular dendritic cells (FDCs) in LZ. Low-affinity B cells will undergo apoptosis, whereas high-affinity B cells receive help from T follicular helper (T_FH_) cells to undergo CSR, and subsequently either re-enter the SHM‒proliferation cycle in the DZ or exit the GC as plasma cells or memory B cells. AIRE in B cells limits autoantibody generation by restraining excessive AID activity in the GC.

## DISCUSSION

### AIRE as A Paradigm of B Cell-Intrinsic Intracellular Immune Checkpoint

The concept and importance of immune checkpoints are now widely appreciated by scientists and clinicians. Immune checkpoints refer to molecules that mediate negative regulation of immune responses and therefore have a crucial role in maintaining immunological self-tolerance and avoiding autoimmunity. Many checkpoint molecules can be hijacked by tumors or pathogens for immune evasion and have therefore emerged as therapeutic targets for cancer and infectious diseases. To date, the majority of immune checkpoints under research investigation or clinical development encompass cell surface ligands and receptors that are involved in antigen-presenting cell (APC)–T cell interactions in immune-inductive sites (i.e., secondary lymphoid organs) or immune-effector sites (Sharma and Allison, 2015). Here, we demonstrate AIRE expression in the GC stage of B cell differentiation and its negative regulation on AID-mediated peripheral antibody diversification. Consequently, AIRE deficiency in B cells results in exaggerated antibody diversification and autoimmunity. Importantly, increased SHM of *Aire*^−/−^ donor B cells was only seen in class-switched isotypes but not the IgM or IgD compartment, indicating an effect of AIRE only in AID-experienced B cells. This is not only consistent with the GC-specific AIRE expression in peripheral B cells but also suggests that the function of AIRE described here is not a secondary effect of thymic B cells. The result from *Aire*^+/+^ and *Aire*^−/−^ B cell chimeras further corroborates this conclusion and confirms the B cell-intrinsic role of AIRE in limiting antibody diversification. Therefore, AIRE represents a paradigmatic B cell-intrinsic intracellular immune checkpoint of humoral immunity (**Figure 7I**).

### Functionally Significant AIRE Expression in Broader Cell Types

Besides mTECs (Anderson et al., 2002), AIRE protein expression has been previously reported in thymic B cells as well as several normal or malignant cell types of the hematopoietic or non-hematopoietic lineage in the periphery in humans and/or mice (Bianchi et al., 2016; Gardner et al., 2008; Gardner et al., 2013; Hobbs et al., 2015; Lindh et al., 2008; Poliani et al., 2010; Yamano et al., 2015). Intriguingly, little or no AIRE protein was seen in mouse GC B cells in a previous study, which was thought to result from B cell receptor (BCR)-mediated inhibition of AIRE induction; however, mouse B cells still expressed markedly elevated levels (∼20 fold) of AIRE transcript when they were stimulated with anti-CD40 in the presence of anti-IgM (Yamano et al., 2015). In contrast, we present evidence obtained using multiple approaches to demonstrate AIRE protein expression in human and mouse GC B cells *in vivo*, and have also found much higher levels of AIRE transcript in mouse GC B cells than in non-GC B cells (**Figure 1H**). Therefore, BCR triggering does not completely abolish AIRE induction in GCs under physiological conditions. Consistent with our finding, BCR signaling in the GCs is reduced due to high phosphatase activity (Khalil et al., 2012) and, in some cases, BCR specificity may not be required for entry into GCs (Silver et al., 2018). It is additionally unclear whether AIRE is temporally regulated during B cell activation. Our data indicate an impact of AIRE on both CSR, which occurs soon after antigen exposure prior to GC formation (Roco et al., 2019), as well as SHM, which is largely carried out within the GC. AIRE is likely expressed in both the early and late phases of the GC reaction, though whether its expression is dynamically regulated within the GC is still unknown. Nonetheless, the role of AIRE in inhibiting AID shown here is in line with BCR engagement in facilitating CSR and SHM of antigen-specific B cell clones and their affinity maturation by downregulating AIRE.

Although this study focused on the function of AIRE in the regulation of antibody diversification, it remains unknown whether AIRE in GC B cells has a significant impact on TSA expression and Treg induction as seen in mTECs and thymic B cells (Anderson et al., 2002; Malchow et al., 2013; Yamano et al., 2015), or an impact on CD4^+^ T cell inactivation and selection similar to eTACs in secondary lymphoid organs (Gardner et al., 2008; Gardner et al., 2013). Our data suggest that GC B cell AIRE may not be critical in maintaining Treg cells in secondary lymphoid organs, as we observed comparable T_FR_ responses in immunized μMT recipients of *Aire*^+/+^ and *Aire*^−/−^ donor B cells and similar expression of co-stimulatory and co-inhibitory molecules on these donor B cells. However, consistent with its function as a transcriptional regulator (Fang et al., 2024; Giraud et al., 2012; Oven et al., 2007), we observed a clear interaction between AIRE and pSer5 Pol II (**Figure 6F**). In B cells undergoing CSR and SHM, this interaction is particularly important as a targeting mechanism for AID (Chaudhuri et al., 2003; Pavri et al., 2010). In addition, the CARD domain of AIRE, which is necessary for its interaction with AID (**Figure 5F**), is also critical for AIRE polymerization and the formation of nucleation sites that facilitate positive feedback loops to promote transcription (Huoh et al., 2024), indicating that either these condensates are also critical for preventing AID from being recruited to Pol II or AIRE may perform alternative functions in B cells compared to mTECs. Interestingly, AIRE in is also expressed during spermatogenesis and is a regulator of germ cell apoptosis and gene expression independent of the adaptive immune system (Radhakrishnan et al., 2016; Schaller et al., 2008), highlighting its cell-type-specific function and potential evolution as a master regulator of germ cell development that facilitates epigenetic remodeling to promote genomic stability. Nevertheless, additional comprehensive studies on transcriptional regulation by AIRE in GC B cells may provide further insight into a possible role in regulating the expression of TSAs to promote peripheral T cell tolerance.

### AIRE as a Negative Regulator of AID

Many proteins have been reported to interact with and/or regulate AID in B cells, and they are thought to function at various steps of the molecular cascade of antibody diversification ranging from chromatin remodeling, AID targeting, transcriptional regulation, RNA processing, DNA repair and post-translational protein modification (Casellas et al., 2016; Vaidyanathan et al., 2014; Xu et al., 2012). Among the AID-interacting partners identified, AIRE is one of the few, the deficiency of which enhances AID function, whereas the deficiency of many others impairs AID function. This provides a unique way to up-regulate the activity of AID in B cells. Whether AIRE interacts with AID directly or indirectly via other factors remains to be elucidated. Our results suggest the requirement for the CARD and NLS domains of AIRE in the interaction with AID. CARD is critical for AIRE’s interaction with bromodomain-containing protein 4 (Brd4) which binds acetylated lysines in CARD and bridges AIRE with the positive transcription elongation factor b (P-TEFb) complex to induce ectopic gene expression in mTECs by promoting Pol II elongation (Giraud et al., 2014; Giraud et al., 2012; Oven et al., 2007; Yoshida et al., 2015). The increased interaction of AID with IgH S region and transcriptionally paused Pol II in *Aire*^−/−^ CH12 cells undergoing CSR is therefore consistent with this body of literature, considering AID is targeted to DNA sites of Pol II pausing (Pavri et al., 2010). The genomic uracil dot blot assay we employed directly detects the product of AID’s enzymatic activity, in contrast to conventional methods, such as flow cytometry, ELISA and antibody sequencing, that measure the outcome of CSR and SHM as an indirect readout of AID’s function. Increased generation of genomic uracils in *Aire*^−/−^ CH12 cells undergoing CSR points to the regulation of AID by AIRE at steps upstream of the deamination reaction, which further agrees with the role of AIRE in restraining AID targeting to its DNA substrate. However, the involvement of other mechanisms upstream or downstream of the deamination reaction, such as the regulation of chromatin availability or DNA repair by AIRE (Abramson et al., 2010; Koh et al., 2018), cannot be discounted.

### B Cell-Extrinsic and -Intrinsic AIRE Deficiency May Differentially Contribute to Autoimmunity and Immunodeficiency in APS-1

APS-1 is caused by mutations in a single gene but has emerged as a disease with complex pathogenesis. In fact, it has been classified as a type IV PID, a disease of immune dysregulation (Al-Herz et al., 2011). APS-1-associated CMC amidst the multi-organ autoimmune manifestations indicates the co-occurrence of immunodeficiency and autoimmunity, which are two conditions that are thought to occur usually at the opposite ends of the clinical immune spectrum. Such a seemingly paradoxical feature is now being found in an increasing number of primary and acquired immunodeficiencies and autoimmune diseases, such as selective IgA deficiency (SIgAD), immune dysregulation-polyendocrinopathy-enteropathy-X-linked syndrome (IPEX), severe combined variable immunodeficiency (CVID), acquired immunodeficiency syndrome (AIDS), systemic lupus erythematosus (SLE) and diabetes mellitus (DM) (Bacchetta and Notarangelo, 2013; Grammatikos and Tsokos, 2012; Zandman-Goddard and Shoenfeld, 2002). Our work revealed that, besides the central and peripheral abnormalities in T cell tolerance extrinsic to B cells, APS-1 involves previously unknown B cell-intrinsic dysregulation in peripheral antibody diversification, and this can engender humoral autoimmunity, such as the production of autoreactive antibodies against T_H_17 effector cytokines. Our findings offer a plausible mechanistic explanation to the clinical observation of the presence of high affinity autoreactive neutralizing antibodies in these patients (Kisand et al., 2010; Meyer et al., 2016a; Puel et al., 2010) and the causes of defective anti-*Candida* immune defense, thereby arguing for the concept that the seemingly contradictory autoimmunity and immunodeficiency can go hand-in-hand if there is an overproduction of autoreactive antibodies that impair protective immunity against pathogens. Of note, B cells were required to cause the multi-organ inflammation in *Aire*^−/−^ mice not by producing autoantibodies, but by mediating early T cell priming and expansion (Gavanescu et al., 2008). This suggests that the multi-organ autoimmunity and CMC-associated immunodeficiency in APS-1 are differentially contributed by the B cell-extrinsic and -intrinsic consequences of AIRE deficiency. We think that this scenario, if proven to be true, could be a paradigm applicable to understanding many other immunodeficiencies co-presenting with autoimmunity that involve humoral immune dysfunction. Indeed, an interesting example that mirrors this scenario is seen in AID-deficient individuals and mice that suffer from immunodeficiency due to the lack of CSR and SHM in the periphery (Muramatsu et al., 2000; Revy et al., 2000) but also have B cell autoimmunity likely due to missing AID’s potential function in purging autoreactive immature B cells in the BM (Cantaert et al., 2015; Meyers et al., 2011). In addition, APS-1 patients also present with autoreactive antibodies to other cytokines, such as type I IFNs (Karner et al., 2013; Meyer et al., 2016b), and it is currently unclear whether these autoantibodies are regulated by B cell intrinsic AIRE, as only T_H_17-associated cytokines, which are critical for controlling fungal pathogens, were tested in this study. Further, although we observed AIRE expression in GC B cells from Peyer’s patches (**Figure S1J**), how this mechanism affects the development of IgA-producing B cells and whether this has any impact on gastrointestinal (GI) disorders such as IBD is yet to be determined. Future studies may include additional models such as viral infections, type 1 diabetes, or GI inflammation to shed more light on the extent and impact of autoantibody generation by AIRE-deficient B cells and potentially offer a new direction for developing therapies that specifically and actively target the various aspects of pathogenesis of APS-1 and these other diseases.

### AIRE Ablation as an Approach of Therapeutic Antibody Generation

Besides offering new mechanistic insights into the regulation of the GC antibody diversification machinery and the unique production of high-affinity neutralizing autoantibodies in APS-1, our study underscores the important and emerging idea that immune tolerance mechanisms can be barriers to the generation of effective immunity (Khan et al., 2014; Schroeder et al., 2017), and controlled breaching of peripheral tolerance can permit neutralizing antibody responses that can be therapeutically beneficial. Remarkably, we found increased Ig framework region SHMs in class-switched antigen-specific *Aire*^−/−^ B cells. Such an observation is reminiscent of the hallmarks of broadly and potently neutralizing antibodies against certain pathogens such as HIV-1 (Klein et al., 2013), the generation of which remains one of the most important mysteries as well as challenges in immunology. Our findings therefore point to a new effective strategy for producing such antibodies by removing a critical B cell-intrinsic brake of AID, which is AIRE. We envisage that further mechanistic elucidation of how B cell-intrinsic AIRE regulates AID function could assist in cracking the elusive molecular underpinning of the generation of broadly neutralizing antibodies and lead to substantial advances in antibody-based immunotherapies against devastating diseases, such as AIDS and cancer.

## STAR★METHODS

Detailed methods are provided in the online version of this paper and include the following:

- KEY RESOURCES TABLE
- CONTACT FOR REAGENTS AND RESOURCE SHARING
- EXPERIMENTAL MODEL AND SUBJECT DETAILS

- Human subjects
- Mice
- Primary cell cultures
- Cell lines
- Microbe strain
- METHOD DETAILS

- Human blood and tissue sample processing and cell isolation
- Mouse blood and tissue cell isolation
- Mouse immunization
- BM and B cell chimeras
- Discrimination of intravascular and tissue leukocytes
- Culture and stimulation or primary B cells
- Generation and validation of *Aire*^−/−^ CH12 cells
- Plasmids
- Transfection
- *C. albicans* culture
- Cutaneous *C. albicans* infection
- Immunoprecipitation
- RNA extraction and quantitative real-time polymerase chain reaction
- Chromatin immunoprecipitation and quantitative real-time PCR
- Protein extraction and Western Blot
- Genomic uracil quantitation
- Conventional flow cytometry
- Imaging flow cytometry
- Immunofluorescence analysis
- ELISA
- IgHV repertoire and mutation analysis
- QUANTIFICATION AND STATISTICAL ANALYSIS

- Statistical analyses
- DATA AND SOFTWARE AVAILABILITY

- Software availability

## AUTHOR CONTRIBUTIONS

J.Z.Z., B.P., and G.W.S. designed and performed research and analyzed data. M.D.P., W.Z., X.L., F.Y., K.C.H., C.-X. L., M.L.W.P., S.W., S.Z., and L.A.P. performed research. J.M.P., A.R., K.Kr., C.C.-R., and A.C. provided specimens and clinical insights and discussed data. N.Y., A.S.B., K.Y., P.P., K.Ki., and B.Q.V. provided reagents and discussed data. B.H. and K.C. designed and directed research, discussed and analyzed data. K.C. conceived the study and wrote the manuscript.

## ACKNOWLEDGMENTS

We are grateful to D. Mathis and C. Benoist (Harvard Medical School, USA) for *AIRE* cDNA plasmids, T. Honjo (Kyoto University, Japan) for *Aicda*^−/−^ mice and CH12 cells, M. Anderson (University of California-San Francisco, USA) for *Aire^Adig^*mice and discussions, T. Hagen (National University of Singapore) for the pcDNA3-eGFP vector, J.X. Ang, G. Lee and Q. Peng (Republic Polytechnic, Singapore) for assistance in creating and validating *Aire*^‒/‒^ CH12 cells, C. Lim (Republic Polytechnic, Singapore) for assistance in constructing *Aire* mutant plasmids, E. van Buren and J. Back (Microscopy and Flow Cytometry Core of the Barbara Ann Karmanos Cancer Institute, USA) for assistance in cell sorting and imaging flow cytometry, and S. Dzinic, K. White and J. Kushner (Animal Model and Therapeutics Evaluation Core of the Barbara Ann Karmanos Cancer Institute, USA) for assistance in some animal experiments. This work was supported by the US National Institutes of Health (R21AI122256 and U01AI95776IOF to K.C.; R01GM057200 to A.S.B.), Shenzhen Medical Research Fund (B2302008 to B.H.), National Natural Science Foundation of China (32370982 to B.H.), Guangdong Science and Technology Department grant (2024B1212030002 to B.H.), Wayne State University (Bridge award to A.S.B.) and a Lee Kuan Yew Postdoctoral Fellowship (to S.Z.).

## SUPPLEMENTAL INFORMATION

### SUPPLEMENTAL TABLES

**Table S1.**
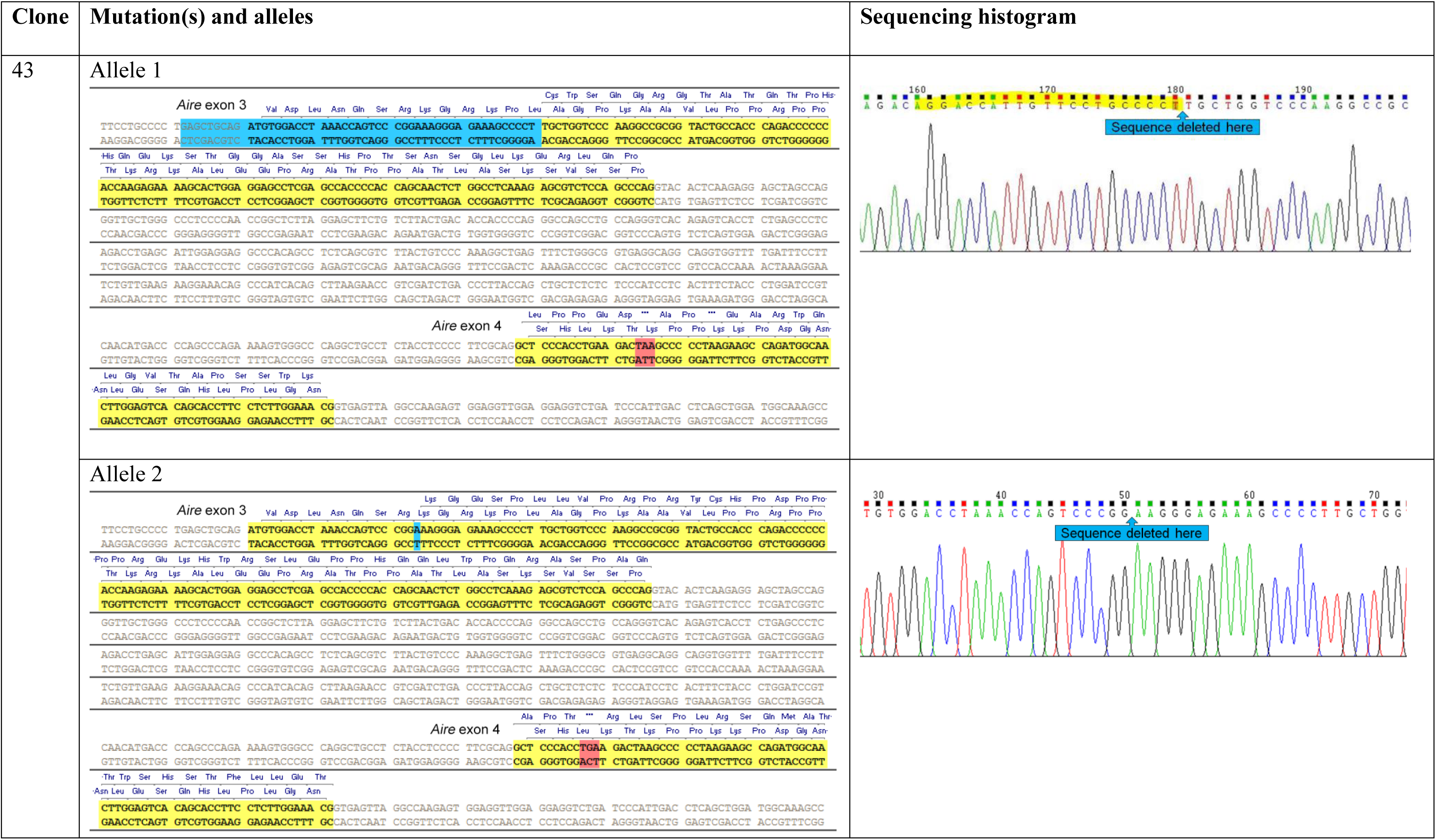
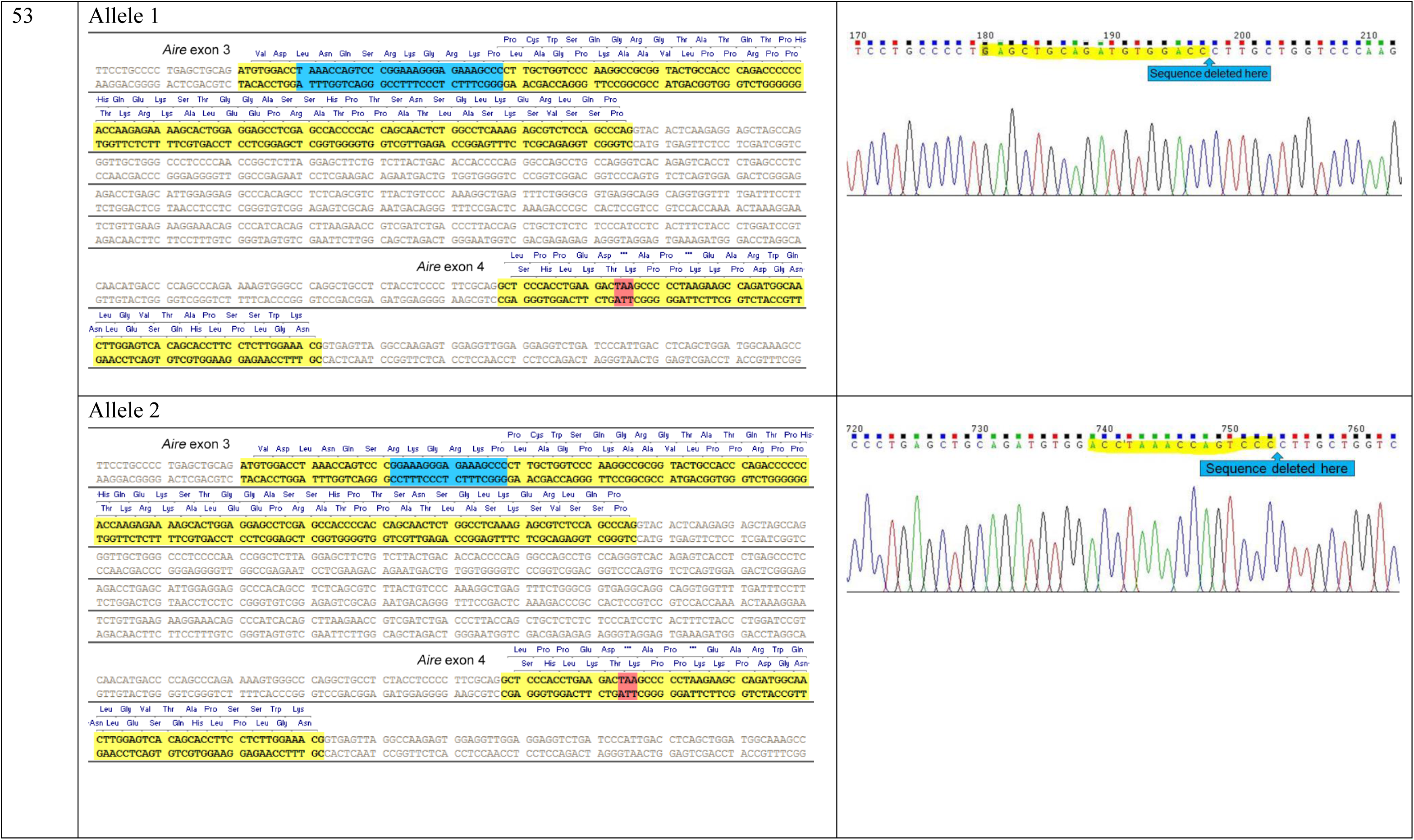
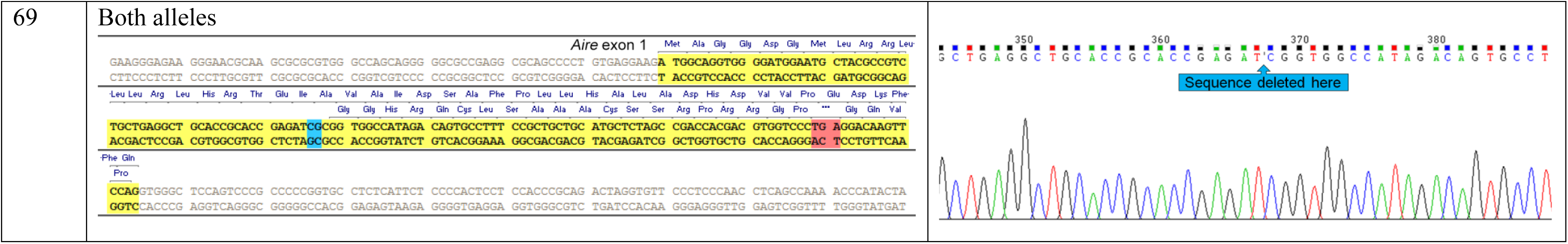
*Aire*^−/−^ CH12 Cell Clones, Related to Figure 4 and Figure 5.

**Table S2.** Human AIRE and AID Constructs, Related to Figure 4 and Figure 5.

**(A) Human AIRE constructs in pcDNA3.1(-)/Myc-His**
| Name | Description | Remaining region | MW with tag (kDa) |
| --- | --- | --- | --- |
| WT | Full length | 1-545 | 60.7 |
| M1 | ΔPHD2 | 1-430 | 48.8 |
| M2 | ΔPHD1, PHD2 | 1-298 | 35 |
| M3 | ΔSAND, PHD1, PHD2 | 1-181 | 23 |
| M4 | ΔCARD | 101-545 | 49 |
| M5 | ΔCARD, ΔNLS | 181-545 | 41 |
| M6 | ΔNLS | 1-100, 181-545 | 52.5 |
| M7 | NLS only | 101-181 | 12 |

**(B) Human AID constructs in pFLAG-CMV2**
| Name | Description | Remaining region | MW with tag (kDa) |
| --- | --- | --- | --- |
| WT | Full length | 1-198 | 26.3 |
| M1 | E58A | 1-198 | 26.3 |
| M2 | NLS of AID replaced with NLS of nucleoplasmin | 1-198 | 26.3 |
| M3 | ΔCatalytic domain | 1-54, 95-198 | 21.4 |
| M4 | ΔAPOBEC-like and ΔNES domains | 1-94 | 13.8 |
| M5 | ΔNLS | 1-8, 27-198 | 24 |
| M6 | G23S | 1-198 | 26.3 |

**Table S3.**
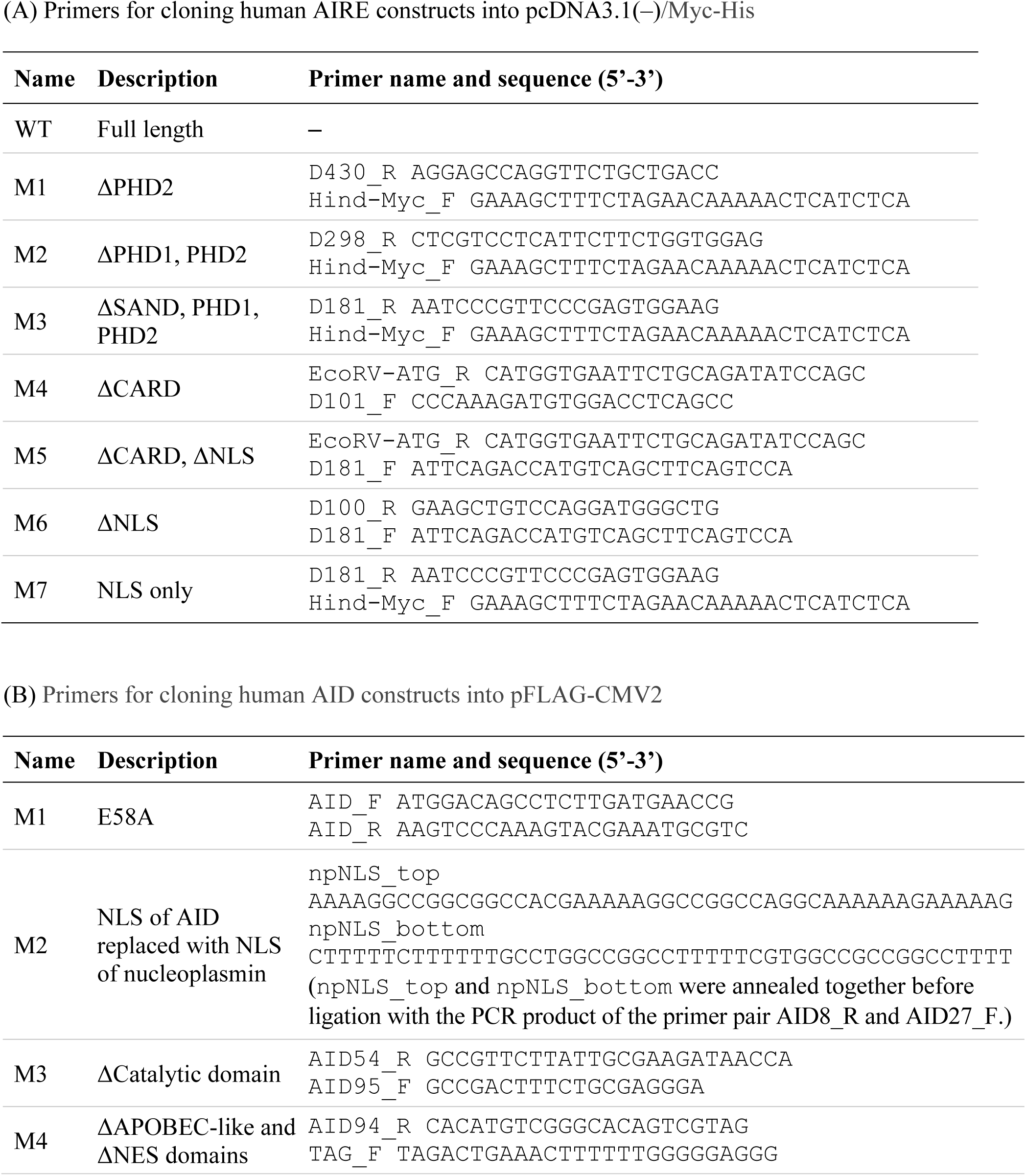
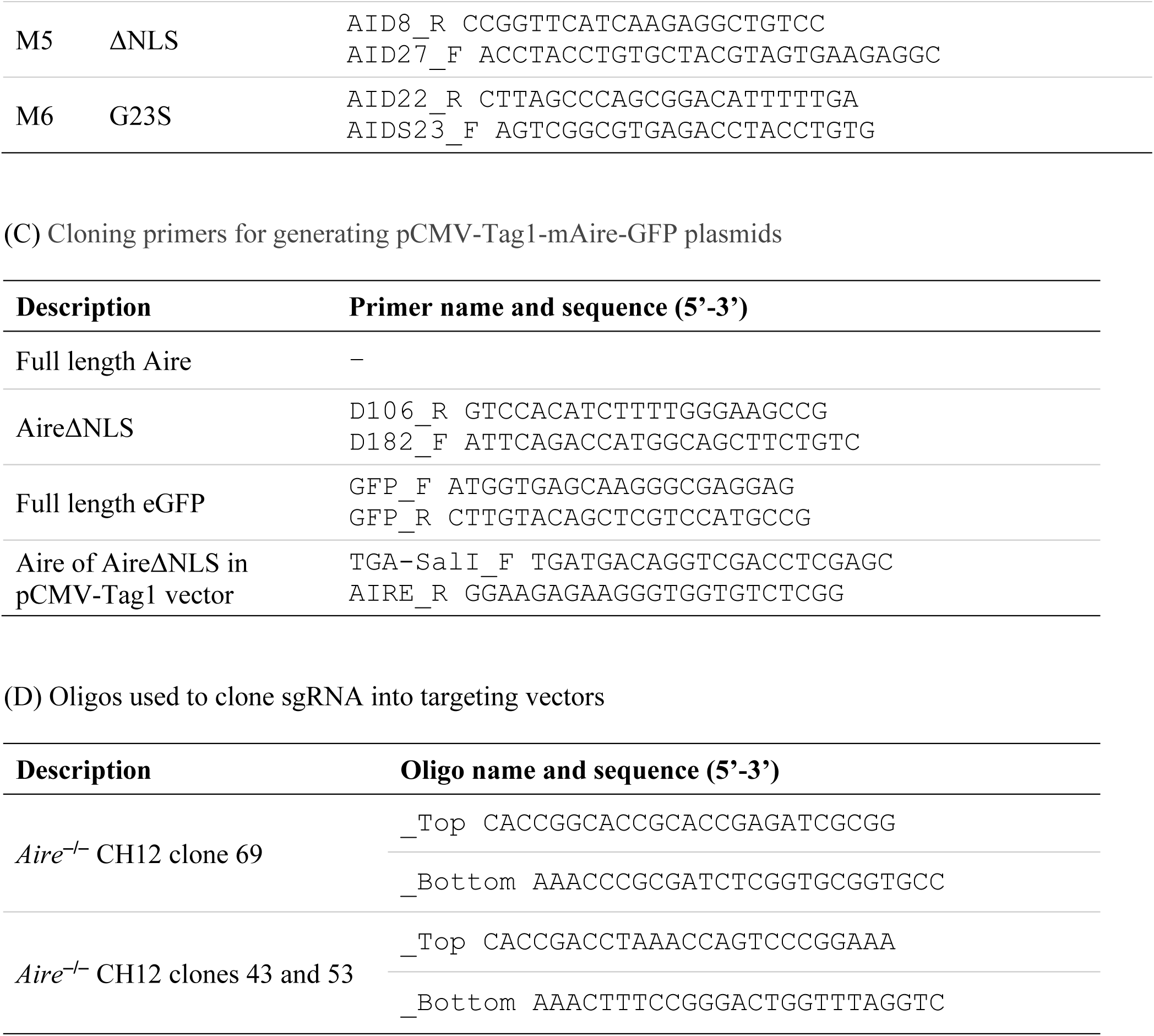
Cloning Primers Used to Generate *Aire*^−/−^ CH12 Clones and AIRE and AID Mutant Molecules, Related to Figure 4 and Figure 5.

### SUPPLEMENTAL FIGURES AND LEGENDS

**Figure S1.**
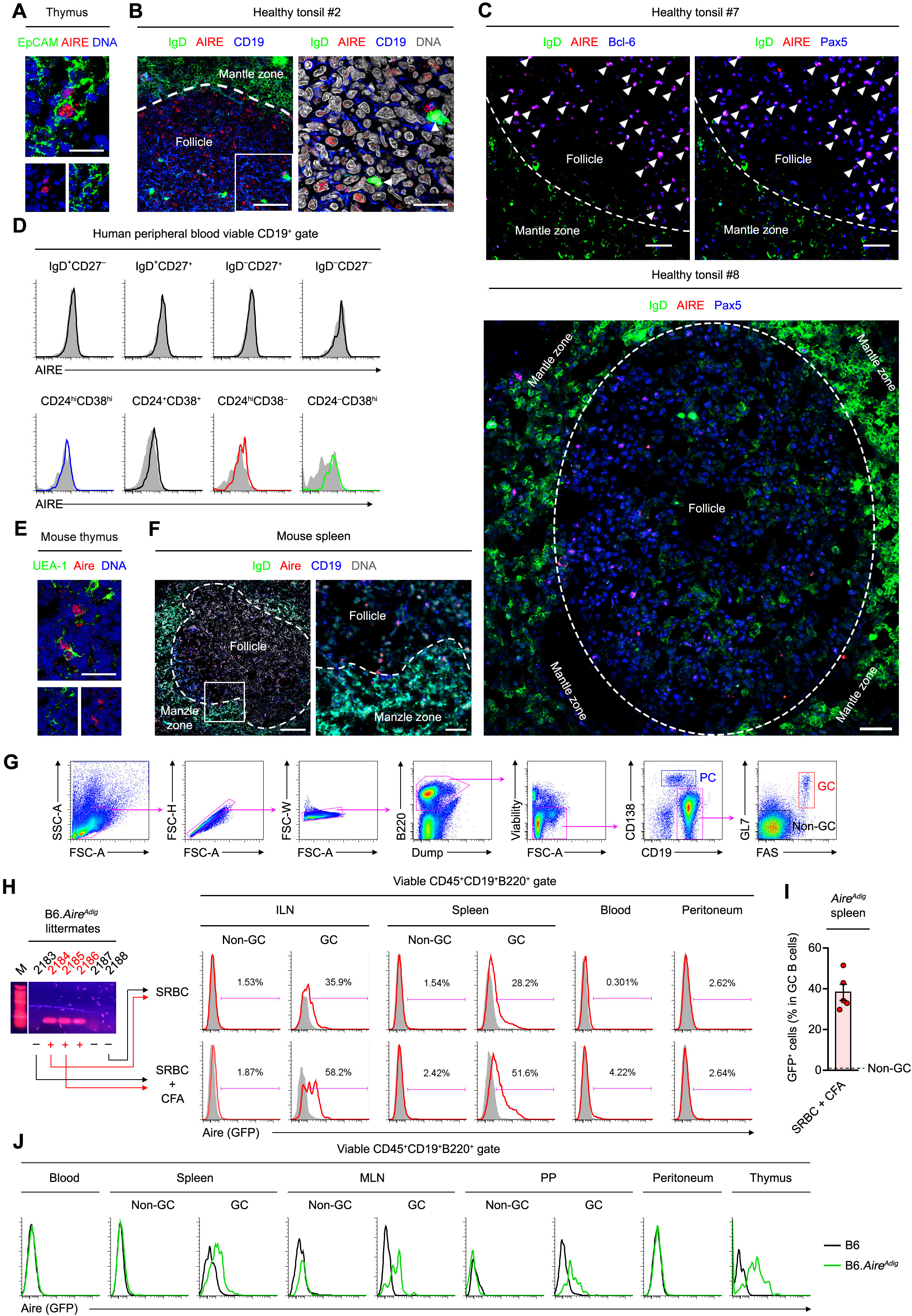
AIRE Is Expressed Specifically in GC B Cells of Human and Mouse Secondary Lymphoid Organs, Related to Figure 1. (A) Immunofluorescence analysis of the thymus of a healthy donor for EpCAM, AIRE and DNA. Bars: 20 μm. (B and C) Immunofluorescence analysis of tonsillar tissues of healthy donors for IgD, AIRE, CD19, Pax5, Bcl-6 and DNA. The dotted lines mark the boundary between tonsil follicular mantle zone and the follicle. Arrow heads point to follicular IgD^+^ plasmablasts (B), Pax5^+^Bcl-6^+^AIRE^+^ GC B cells (C top), and Pax5^+^AIRE^+^ GC B cells (C bottom). Bars: 15 μm (B) and 30 μm (C). (D) Flow cytometric analysis of AIRE expression in human peripheral blood naive (IgD^+^CD27^−^), MZ (IgD^+^CD27^+^), switched memory (IgD^−^CD27^+^), double-negative (IgD^−^CD27^−^) B cells, and transitional (CD24^hi^CD38^hi^), mature (CD24^int^CD38^int^), memory (CD24^hi^CD38^−^) B cells and plasma cells (CD24^−^CD38^hi^). (E) Immunofluorescence analysis of the thymic tissue of a B6 mouse for UEA-1, AIRE and DNA, and the splenic tissue of a B6 mouse immunized with 3 doses of sheep red blood cells (SRBCs) for IgD, Aire, CD19 and DNA, and Bar: 20 μm. (G) Flow cytometric gating strategy for identifying mouse splenic non-GC (CD19^+^B220^+^GL7^−^FAS^−^), GC (CD19^+^B220^+^GL7^+^FAS^+^) B cells and plasma cells (CD19^lo^B220^lo^CD138^+^). (H) Genotypes and Aire expression in ILN, splenic, peripheral blood and peritoneal B cells of a litter of *Aire^Adig^* mice after 1 dose of i.p. SRBC immunization with or without CFA. (I) Percentage of GFP^+^ B cells (mean ± SEM) in splenic GC B cells of *Aire^Adig^* transgene-positive mice (*n* = 5) after 1 dose of i.p. SRBC immunization with CFA. The dotted line indicates of mean value of GFP^+^ B cells in splenic non-GC B cells of these mice. (J) AIRE expression in mouse peripheral blood, splenic, MLN, PP, peritoneum and thymic B cells of B6.*Aire^Adig^* mice after 1 dose of i.p. SRBC immunization without CFA. The data are representative of 6 B6.*Aire^Adig^* and 6 B6 mice that were age- and sex-matched and housed in the same SPF room.

**Figure S2.**
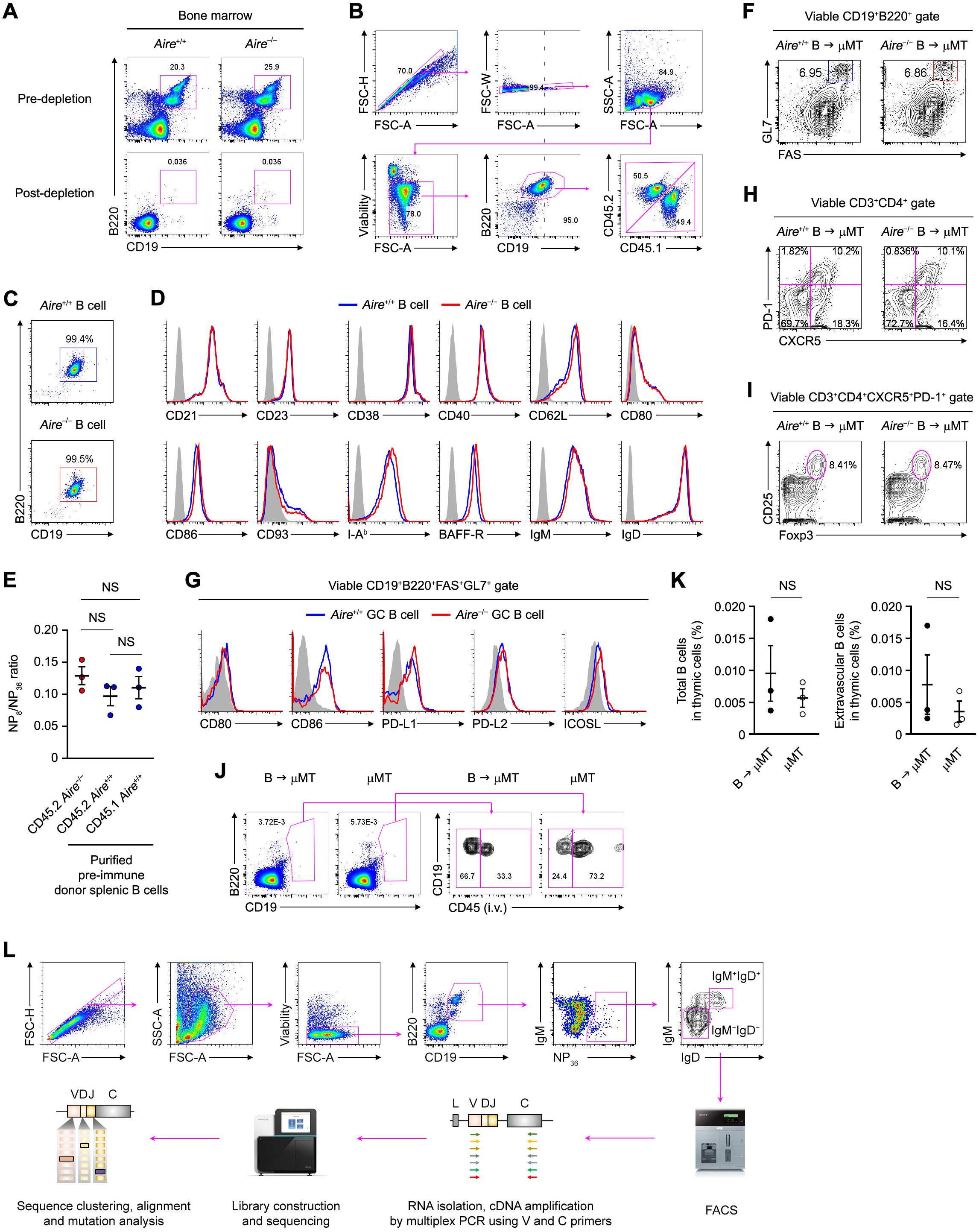
*Aire*^+/+^ and *Aire*^−/−^ Donor BM and B Cells Had a Similar Phenotype Before Transfer, Related to Figure 3. (A) Flow cytometric analysis of CD45.1^+^ *Aire*^+/+^ and CD45.2^+^ *Aire*^−/−^ donor BM before and after B220 cell depletion. (B) Flow cytometry analysis of splenic naive resting B cells that were purified from the spleens of primary μMT chimeras of CD45.1^+^ *Aire*^+/+^ and CD45.2^+^ *Aire*^−/−^ BM and used as donor B cells for the secondary μMT chimeric hosts. The ratio of CD45.2^+^ *Aire*^+/+^ and CD45.2^+^ *Aire*^−/−^ splenic B cells were adjusted to be 1:1 prior to the secondary transfer. (C) Representative purity of CD45.2^+^ *Aire*^+/+^ and CD45.2^+^ *Aire*^−/−^ littermate donor B cells before adoptive transfer into μMT hosts. (D) Cell surface expression of CD21, CD23, CD38, CD40, CD62L, CD80, CD86, CD93, I-A^b^, BAFF-R and IgM and IgD on purified CD45.2^+^ *Aire*^+/+^ and CD45.2^+^ *Aire*^−/−^ littermate donor B cells before adoptive transfer, as determined by flow cytometry. (E) NP_8_-to-NP_36_ binding ratios (mean ± SEM) of pre-immune splenic naive resting donor B cells of CD45.2^+^ *Aire*^−/−^, CD45.2^+^ *Aire*^+/+^ and CD45.1^+^ *Aire*^+/+^ mice, by 1-way ANOVA with Tukey’s post hoc test. (F) Percentage of GL7^+^FAS^+^ GC B cells in the spleens of μMT recipients of either *Aire*^+/+^ or *Aire*^−/−^ B cells that were immunized i.p. with NP_32_-KLH. Flow cytometry was performed 4 d after the last immunization. (G) Cell surface expression of the co-stimulatory or co-inhibitory molecules CD80, CD86, PD-L1, PD-L2 and ICOSL on GL7^+^FAS^+^ GC B cells in the spleens of μMT recipients after immunizations. Shaded histograms indicate the staining using isotype-matched control antibodies. (H and I) Percentage of splenic PD-1^+^CXCR5^+^ T_FH_ cells and PD-1^+^CXCR5^+^Foxp3^+^CD25^+^ T_FR_ cells in the spleens of immunized μMT recipients. The results shown represent 4 experiments, each consisting of B cells from 3–5 age- and sex-matched littermate donor mice and 6–8 age- and sex-matched littermate μMT recipient mice. (J and K) Flow cytometric and statistical analyses of the percentages of total and intravascular B cells in thymic cells of μMT mice that received donor B cells after all the immunizations with NP_32_-KLH. Age and sex-matched unimmunized μMT mice were included as controls. The data are represented as mean ± SEM. (L) The sorting and sequencing strategies for *Aire*^+/+^ and *Aire*^‒/‒^ donor B cells in μMT recipients after immunizations with NP_32_-KLH. NP-specific B cells were sorted based on NP_36_ binding.

**Figure S3.**
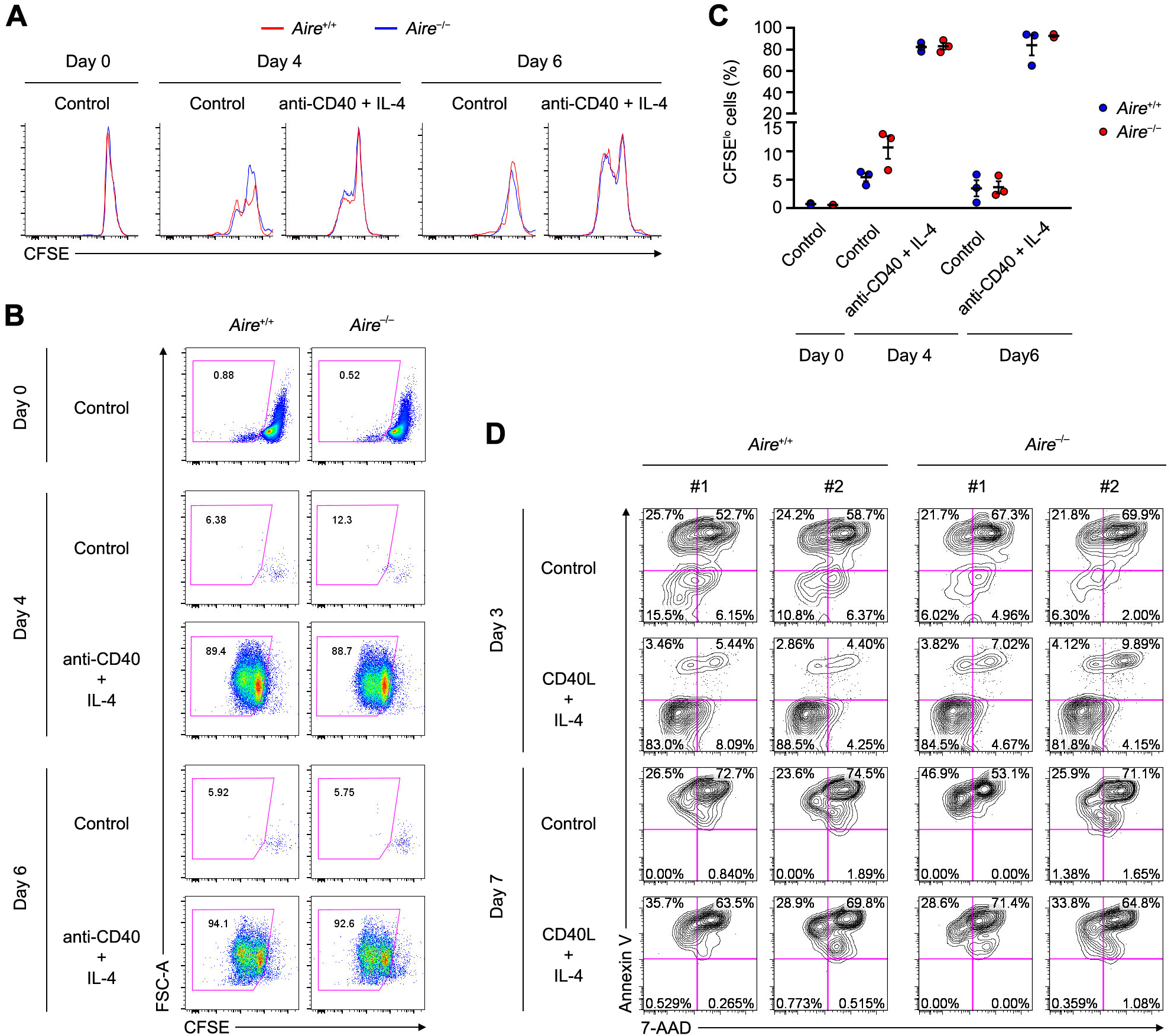
*Aire*^+/+^ and *Aire*^−/−^B Cells Showed Similar Proliferation and Apoptosis *in vitro*, Related to Figure 3. (A and B) CFSE dilution in purified splenic B cells from age- and sex-matched littermate donor *Aire*^+/+^ and *Aire*^−/−^ mice treated with medium (Control) or 5 μg/ml anti-CD40 and 100 ng/ml IL-4 for 4 or 6 d. Non-viable cells were excluded from the analysis. (C) Statistical comparison of the percentage (mean ± SEM) of CFSE^lo^ *Aire*^+/+^ vs. *Aire*^−/−^ splenic B cells (*n* = 3) after 4 or 6 days of stimulation with 5 μg/ml anti-CD40 and 100 ng/ml IL-4, by 2-tailed unpaired *t*-test. The results represent 3 independent experiments. (D) Apoptosis of *Aire*^+/+^ or *Aire*^−/−^ B cells treated with medium (Control) or 500 ng/ml CD40L and 100 ng/ml IL-4 for 3 or 7 d, as determined by Annexin V and 7-AAD staining by flow cytometry. All results shown are representative of 3 experiments, each consisting of cells from 2–3 age- and sex-matched littermate *Aire*^+/+^ and *Aire*^−/−^ mice.

**Fig. S4.**
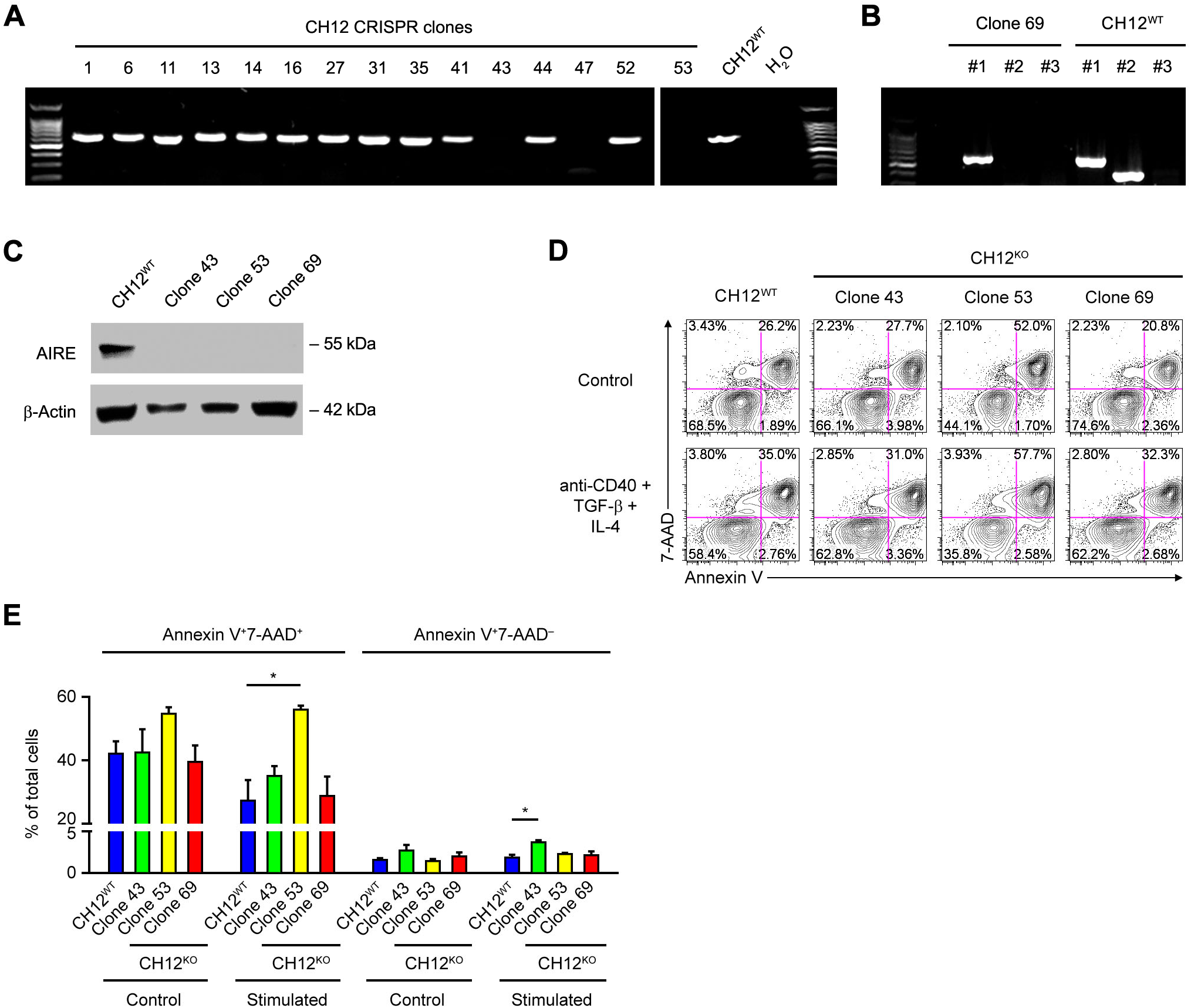
Validation of *Aire*^−/−^ CH12 Cell Clones, Related to Figure 4. (A) Verification of *Aire* mutations in CH12 clones by PCR using primers that only anneal to the WT sequence, giving no amplification in clones 43, 47 and 53. Clone 47 has a 3-bp deletion in both *Aire* alleles causing a single amino acid deletion, and hence was not used in experiments. (B) Verification of *Aire* mutations in both alleles of CH12 clone 69 by PCR showing no amplification using primer pair #2 which anneals to the WT but not the mutated sequence. Primer pair #1 amplifies a sequence immediately downstream of the mutation site, and primer pair #3 is specific for the single-stranded repair template used in CRISPR. (C) Western Blot analysis of AIRE protein expression in WT and *Aire*^−/−^ CH12 cells. (D) Flow cytometric analysis of apoptosis by Annexin V and 7-AAD staining of WT and *Aire*^−/−^ CH12 cells treated with medium (Control) or anti-CD40, TGF-β1 and IL-4 for 3 d. (E) Percentages of late (Annexin V^+^7-AAD^+^) and early (Annexin V^+^7-AAD^−^) apoptotic cells (mean ± SEM) in WT and *Aire*^−/−^ CH12 cells treated with medium (Control) or anti-CD40, TGF-β1 and IL-4 for 3 d. \**P* < 0.05, by 2-tailed *t*-test. The data in D and E represent 4 experiments.

**Fig. S5.**
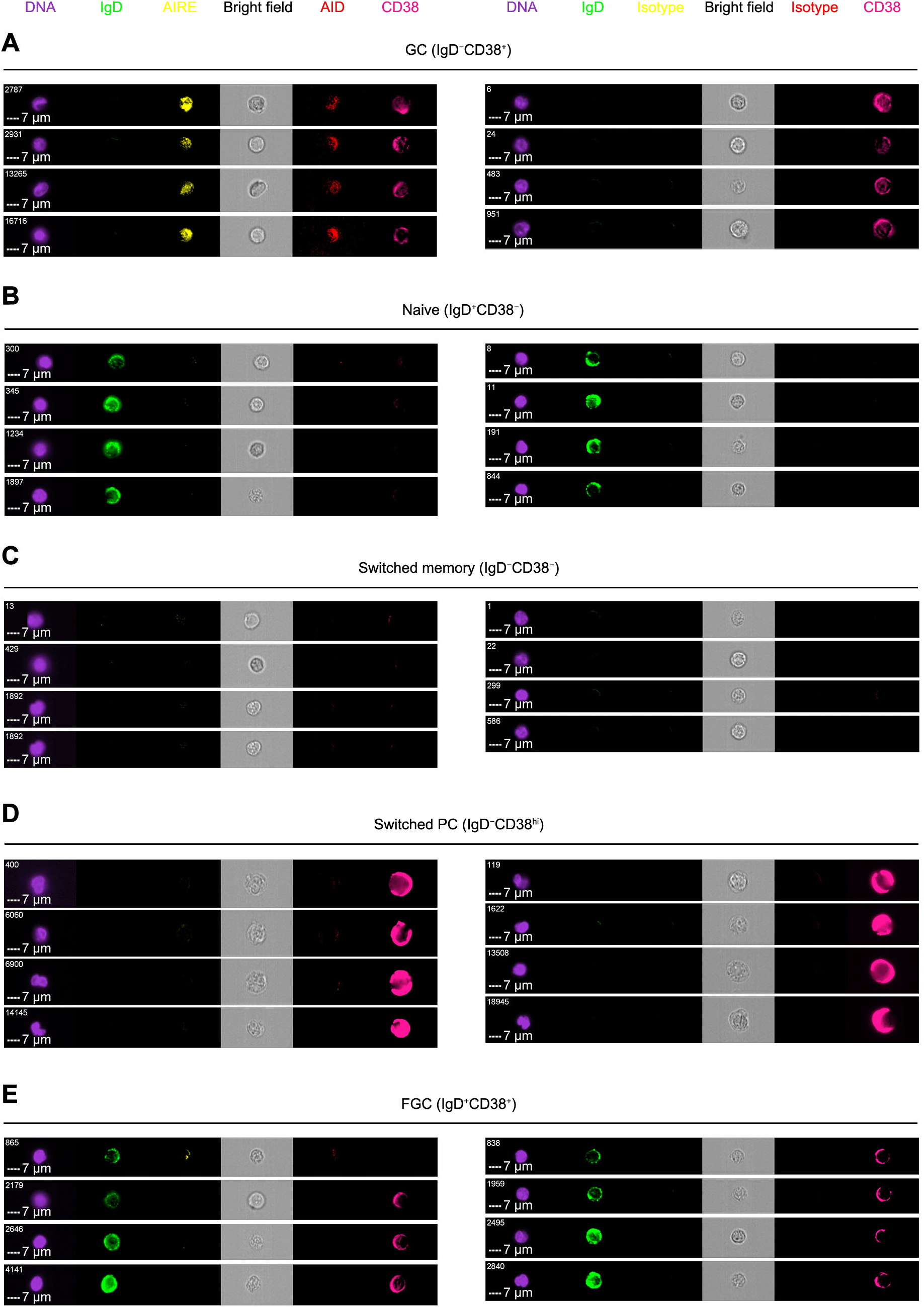
AIRE and AID Co-localize in the Nuclei of GC B Cells, Related to Figure 5. (A–E) Imaging flow cytometry of AIRE and AID in tonsillar IgD^−^CD38^+^ GC, IgD^+^CD38^−^ naive, IgD^−^CD38^−^ switched memory B cells, IgD^−^CD38^hi^ switched PCs and IgD^+^CD38^+^ founder GC (FGC) B cells of a healthy donor. DNA was counter stained with DAPI. Samples stained with isotype-matched control antibodies were used to define the fluorescence baseline for AIRE and AID. Four representative cells in each population stained with AIRE and AID or with isotype control antibodies were shown. Bars: 7 μm.

**Figure S6.**
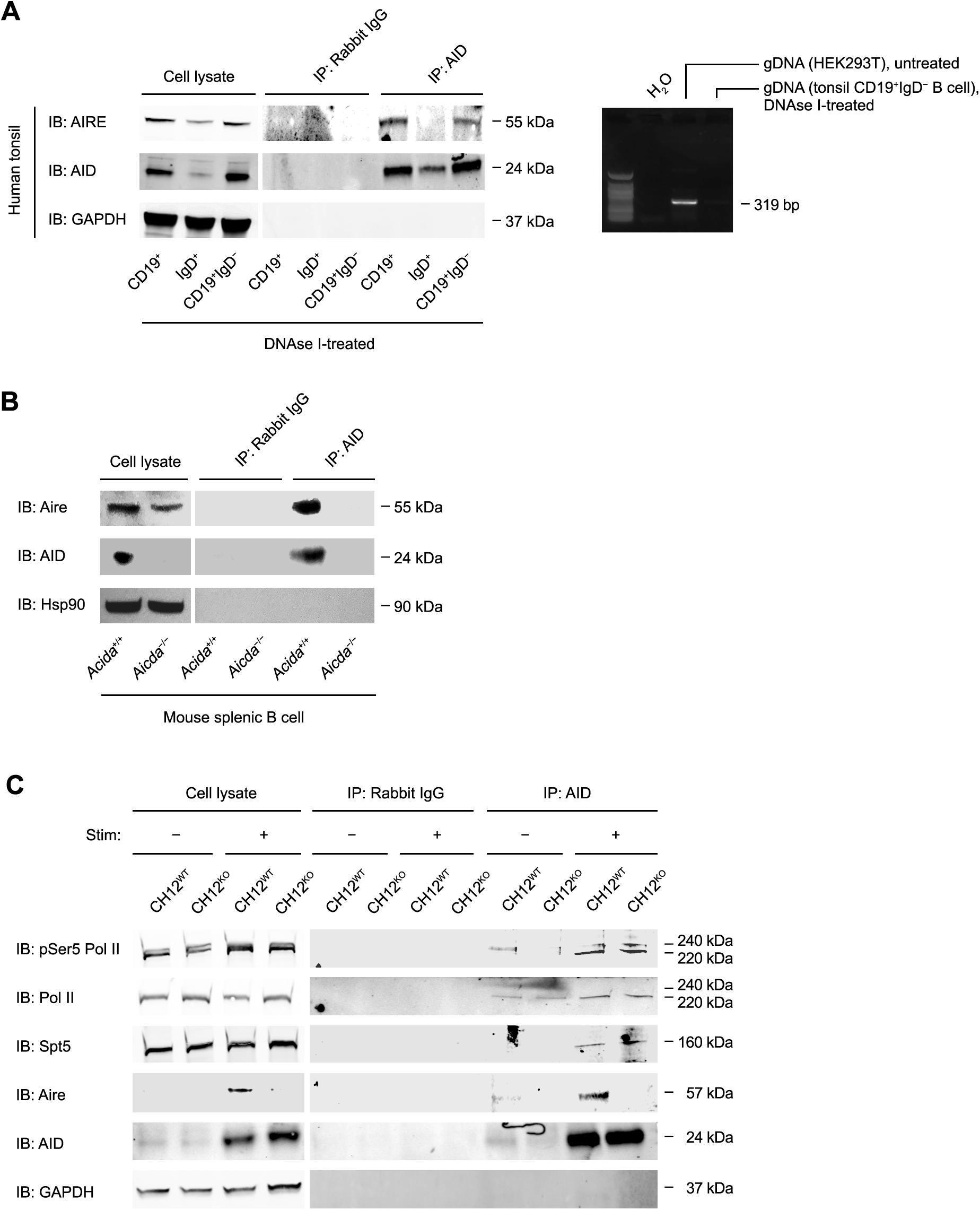
AID Interacts with AIRE in B Cells, Related to Figure 5 and Figure 6. (A) Co-IP of AIRE and AID in tonsillar CD19^+^ total, IgD^+^ naive, and FGC and CD19^+^IgD^‒^ GC and memory B cells of a healthy donor after treatment of the cell lysates with DNAse I. PCR amplification of β-Actin gDNA in DNAse I-treated or untreated cells was also performed. (B) Co-IP of AIRE and AID in splenic B cells of immunized WT or *Aicda*^−/−^ mice. The data represent 2 experiments. (C) Co-IP of AID with pSer5-Pol II, total Pol II, Spt5 and Aire in WT and *Aire*^‒/‒^ CH12 cells after 72 h of treatment without or with anti-CD40, TGF-β and IL-4. The results represent 3 experiments.

**Figure S7.**
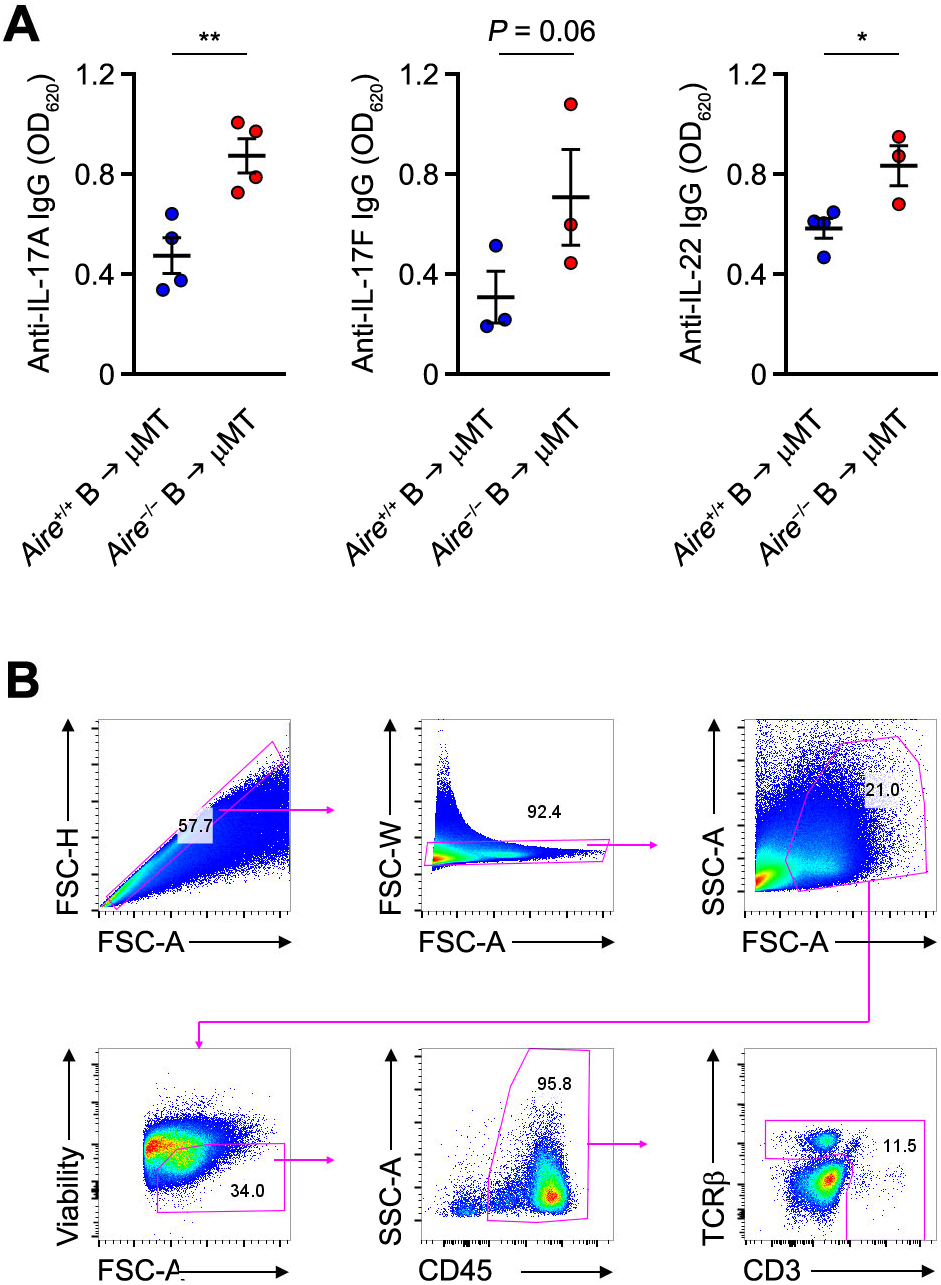
AIRE Deficiency in B Dells Impairs Skin T_H_17 Immunity against *C. albicans*, Related to Figure 7. (A) Levels of autoantibodies (mean ± SEM) binding to IL-17A, IL-17F and IL-22 in the sera of μMT recipient mice of *Aire*^+/+^ or *Aire*^‒/‒^ donor B cells 4 d after infection. \**P* < 0.05, \*\**P* < 0.01, by 1-tailed unpaired *t*-test. (B) Flow cytometric gating strategy for identifying mouse skin viable T cells after *ex vivo* re-stimulation. T cells downregulation CD3 or TCR after *ex vivo* stimulation with PMA and ionomycin; thus CD3^+^ or TCRβ^+^ events were gated for analysis. This gate also included TCRγδ^+^ T cells, which were CD3^+^.

## STAR★METHODS

## KEY RESOURCES TABLE

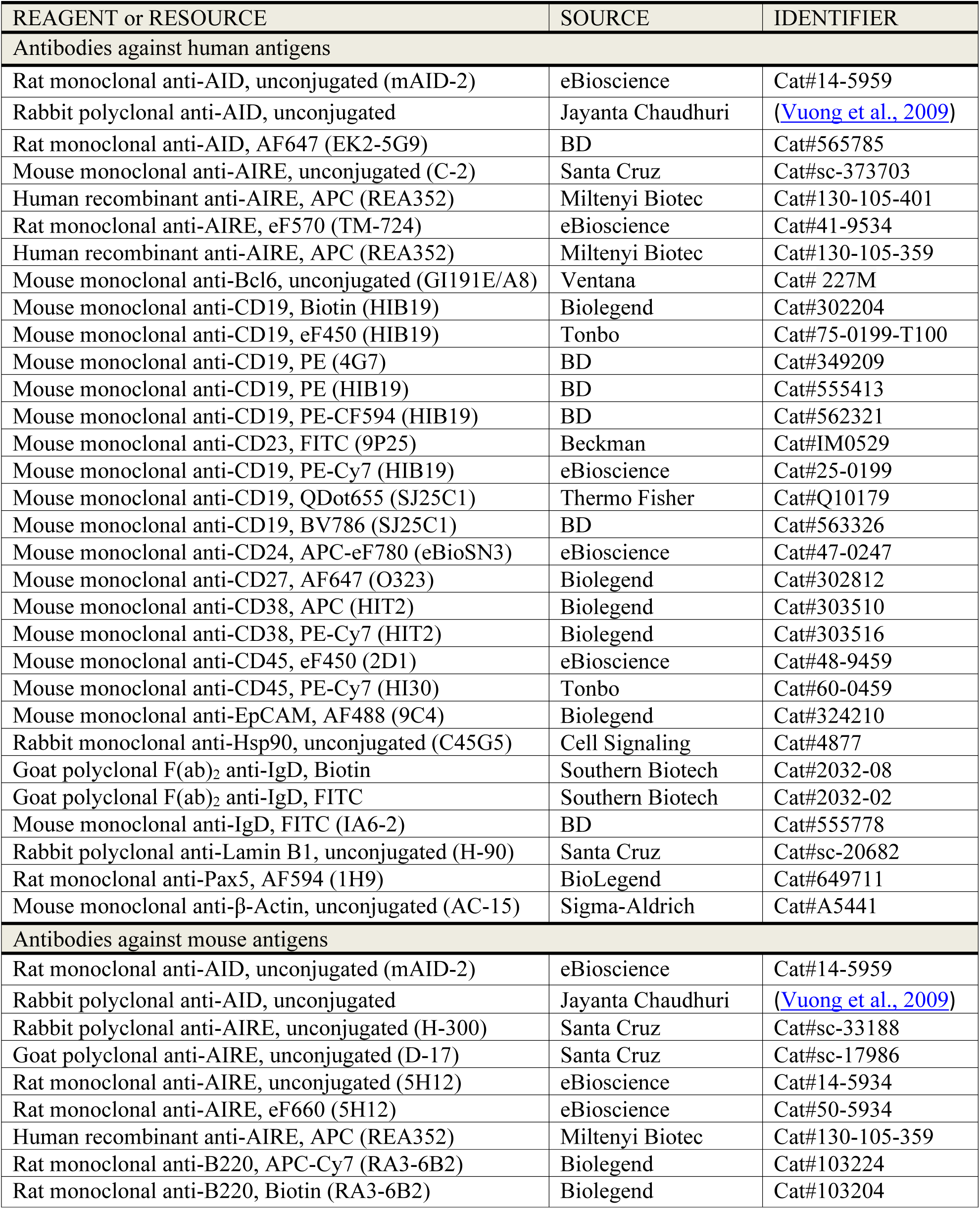
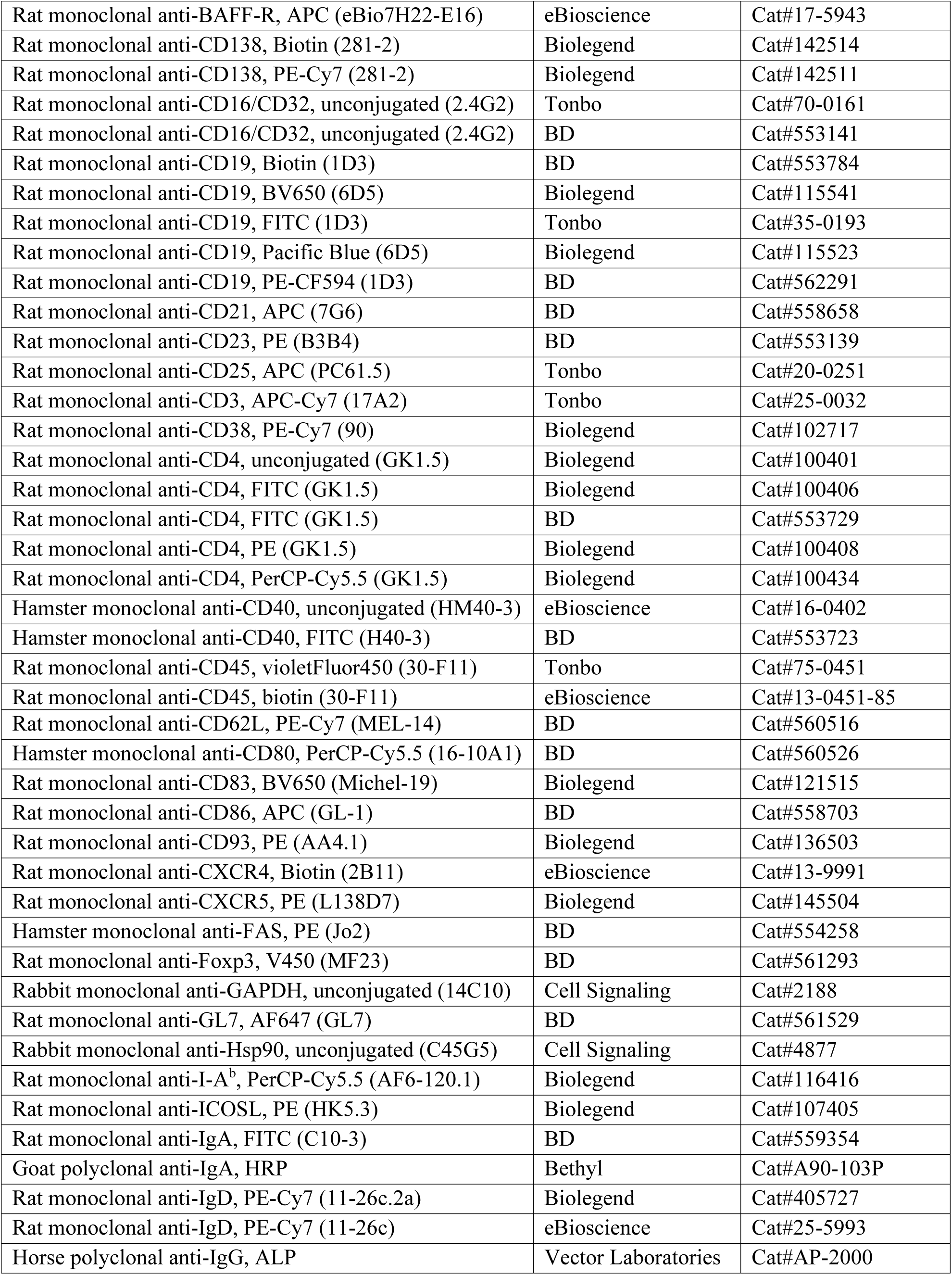
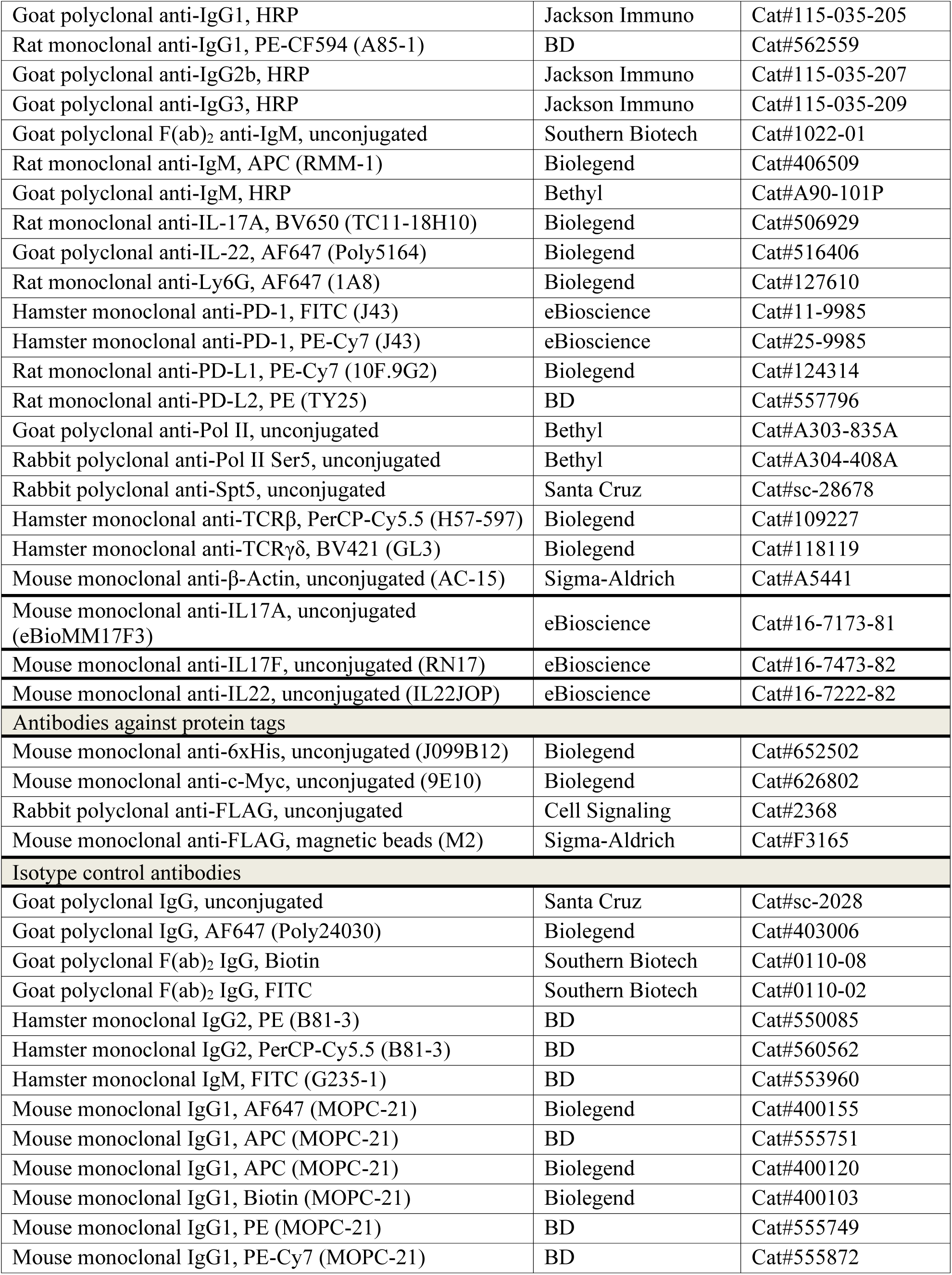
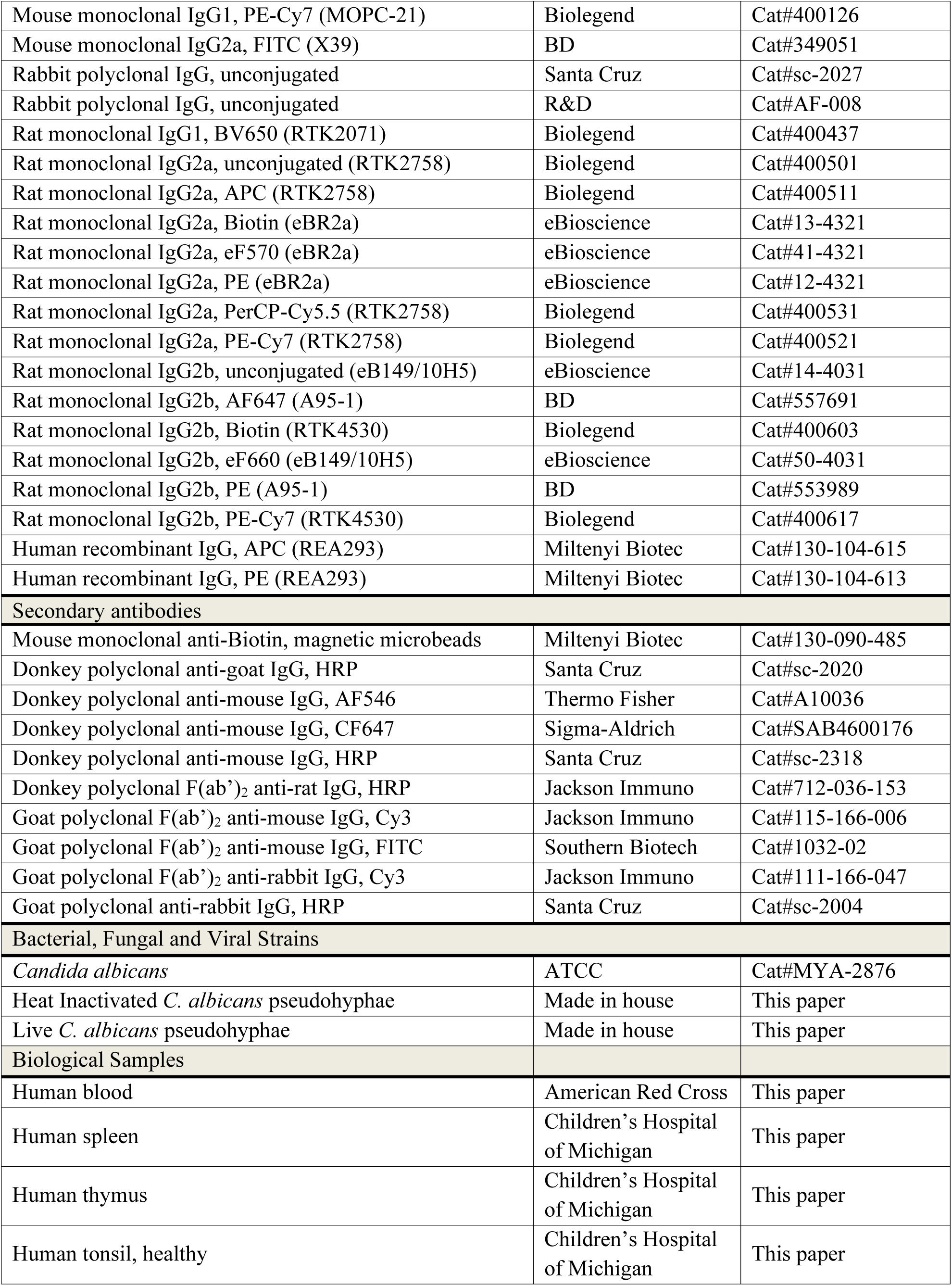
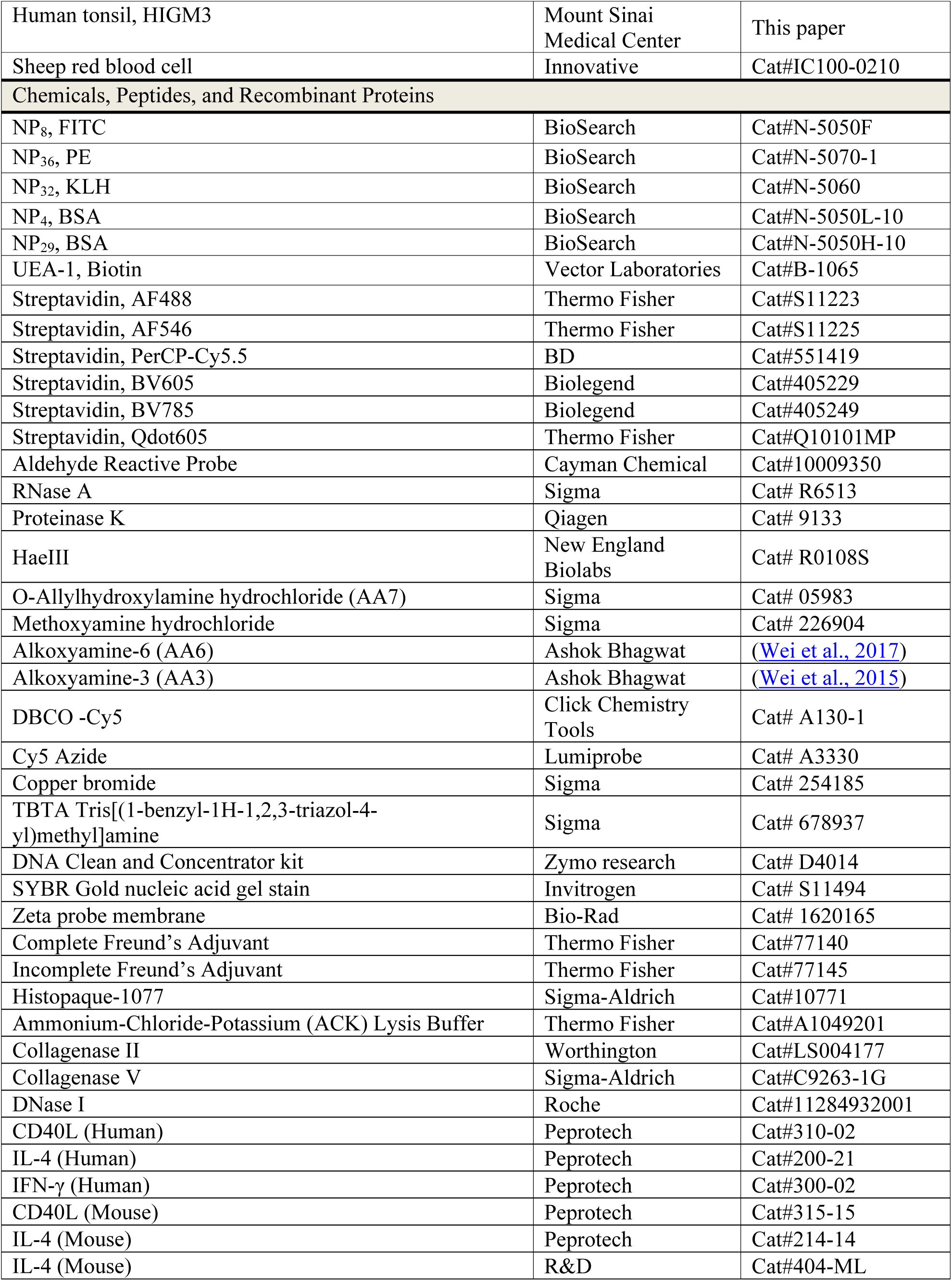
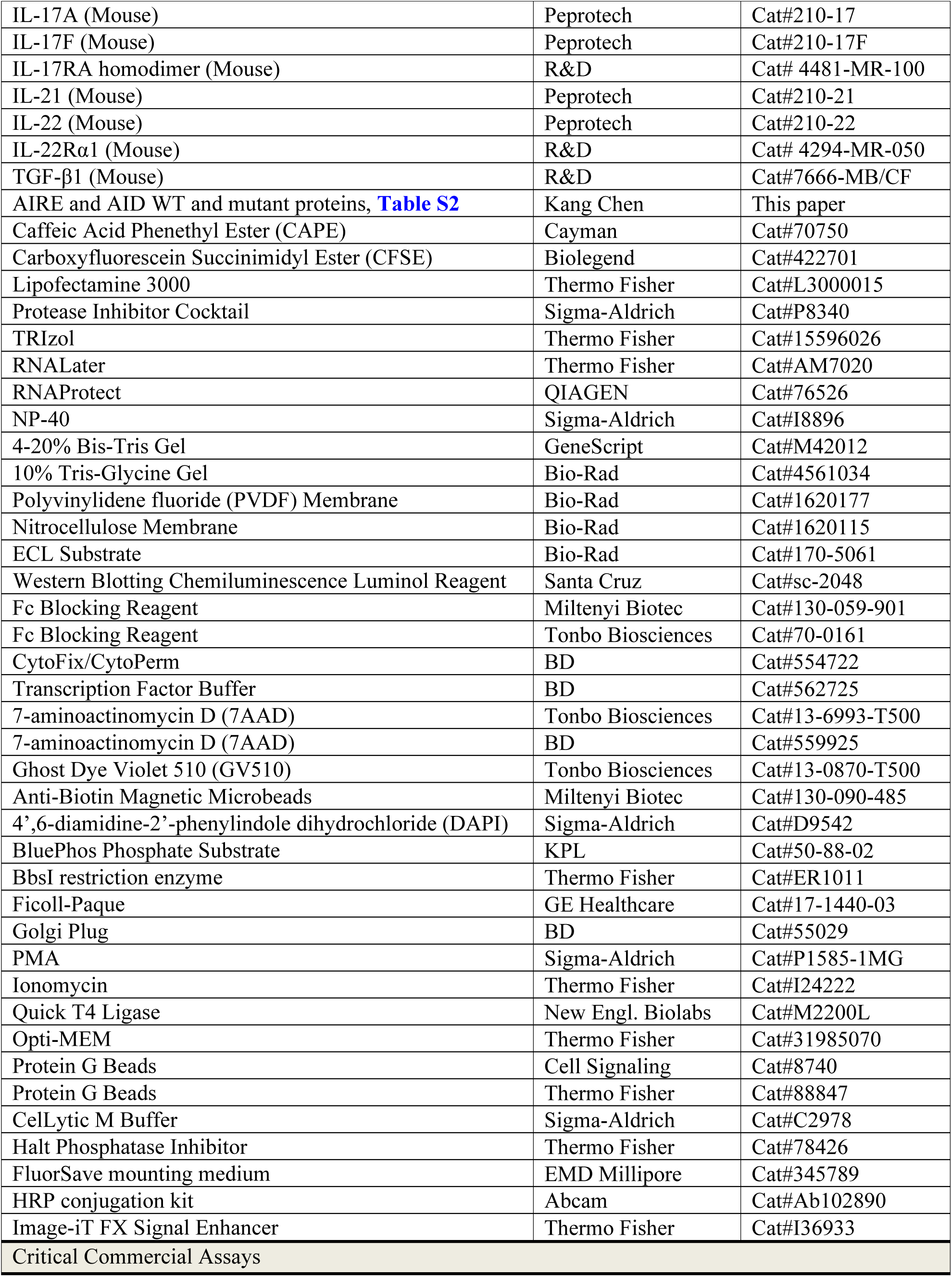
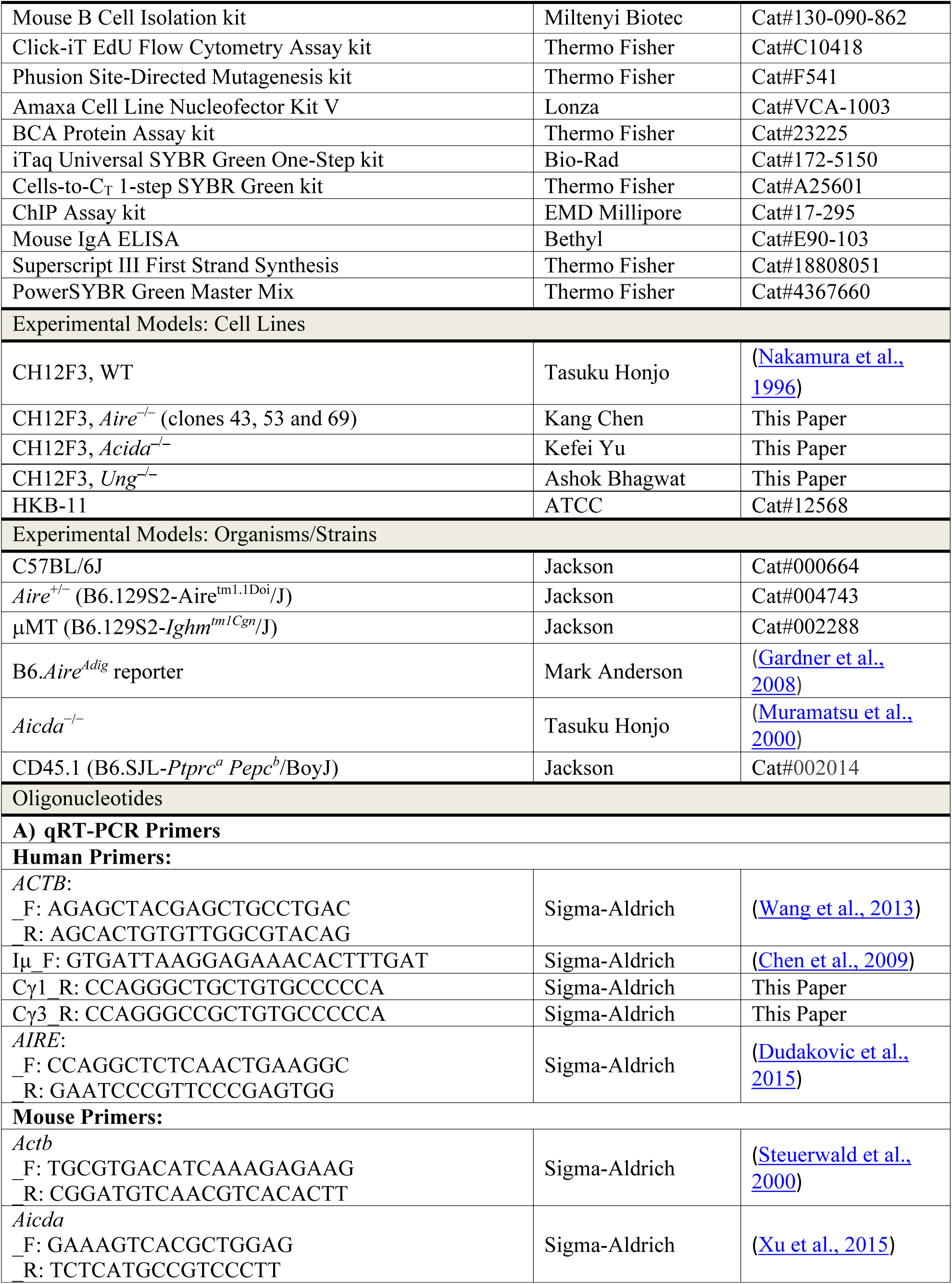
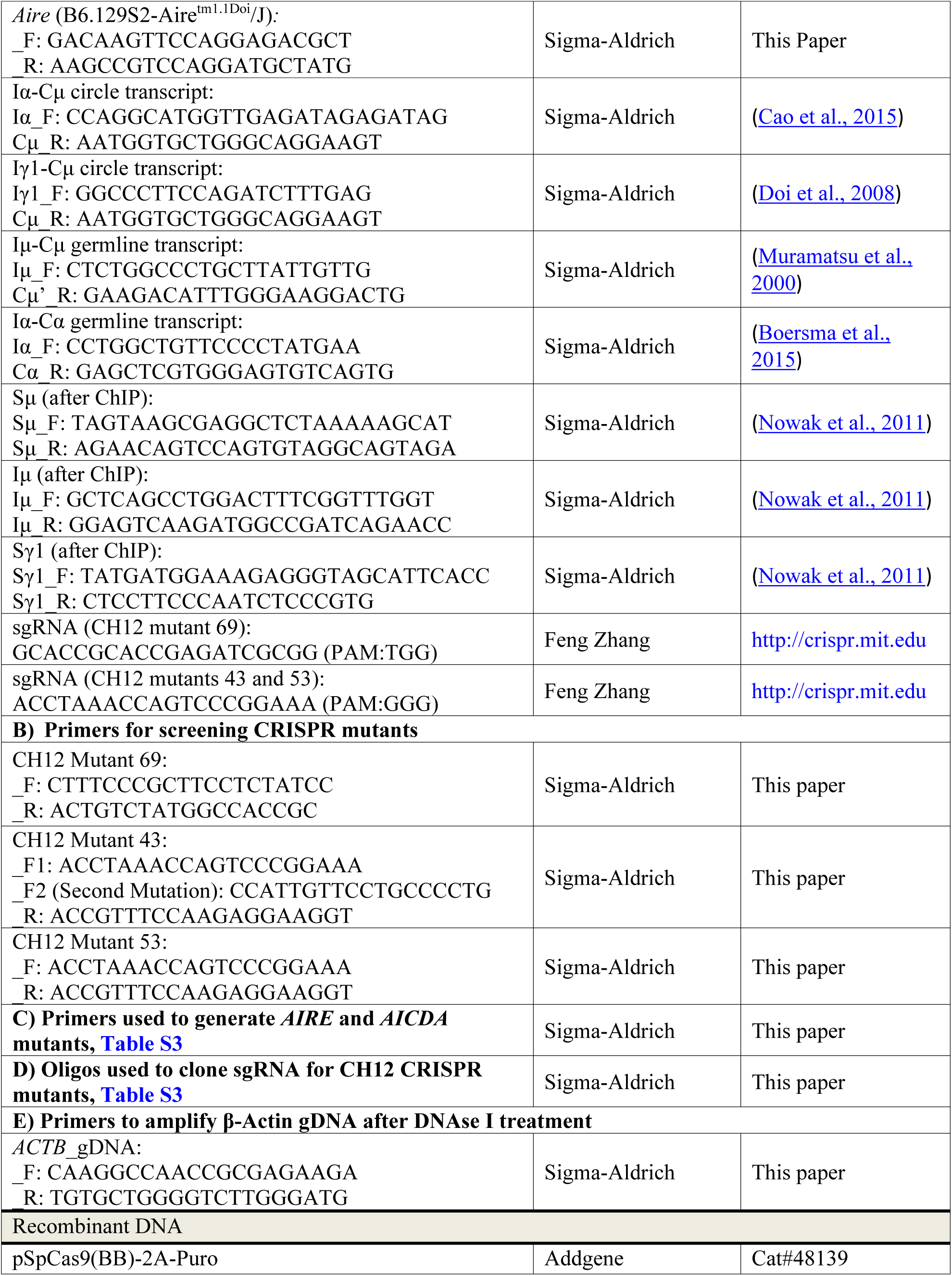
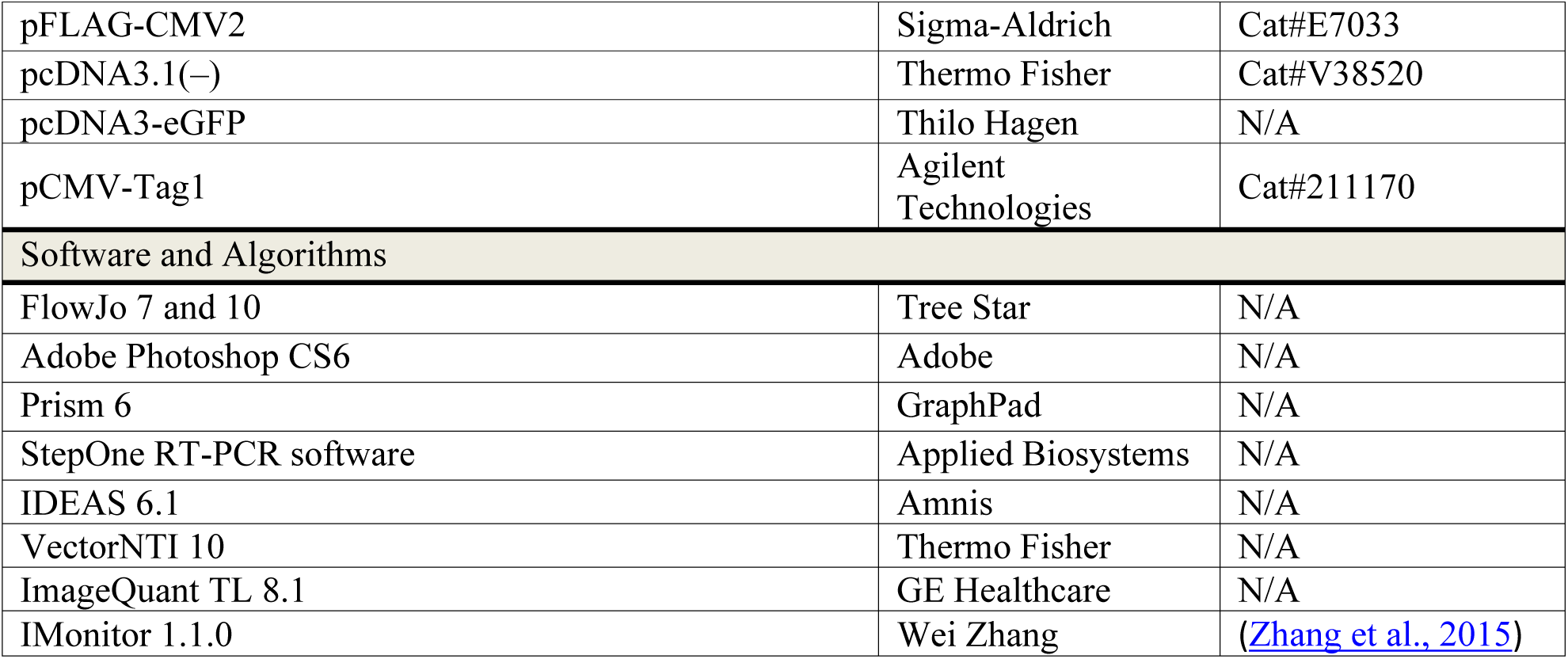

## CONTACT FOR REAGENT AND RESOURCE SHARING

Further information and requests for resources and reagents should be directed to the Lead Contact, Kang Chen.

## EXPERIMENTAL MODEL AND SUBJECT DETAILS

### Human subjects

Autoimmune polyglandular syndrome type 1 (APS-1) patients with loss-of-function mutations in the *AIRE* gene were enrolled in the study with an approved protocol of the Ethics Committee of Medicine of the Hospital District of Helsinki and Uusimaa (HUS), Finland. Hyper-IgM syndrome type 3 (HIGM3) patients with loss-of-function mutations in the *CD40* gene were enrolled in the study with an approved Institutional Review Board (IRB) protocol of the Icahn School of Medicine at Mount Sinai. Peripheral blood leukocytes of anonymous healthy donors were obtained from the American Red Cross with a protocol approved by the IRB of Wayne State University (WSU) and the Detroit Medical Centre (DMC). Tonsil, thymus and spleen tissues were obtained after pediatric tonsillectomy, cardiac surgery and splenectomy, respectively, from the Children’s Hospital of Michigan with an IRB protocol approved by WSU and DMC.

### Mice

C57BL/6J mice, *Aire*^+/−^ and μMT were purchased from the Jackson Laboratory. *Aire^Adig^* mice in C57BL/6 background were previously reported (Gardner et al., 2008). *Aicda*^−/−^ mice (Muramatsu et al., 2000) were generously provided by Dr. Tasuku Honjo (Kyoto University, Japan). These mice were maintained in the same room at the specific pathogen-free (SPF) facility of the Division of Laboratory Animal Resources (DLAR) at Wayne State University. *Aire*^−/−^ mice were generated by mating *Aire*^+/−^ mice. Age- and sex-matched *Aire*^+/+^ littermates were randomly assigned to experimental groups or used as controls for *ex vivo* and *in vivo* experiments. Prior to any experiment, all *Aire* and *Aire^Adig^*mice were genotyped by PCR to ensure the correctness of the genotype. All breeding and experimental protocols were approved by Wayne State University Institutional Animal Care and Use Committee (IACUC).

### Primary cell cultures

Peripheral blood IgD^+^CD27^‒^ naive B cells of healthy subjects or APS-1 patients were sorted and stored in RNALater. Primary blood, spleen and tonsil IgD^+^ B cells were purified by MACS and cultured in RPMI-1640 medium. Mouse cells purified from spleen and lymph nodes were extracted via B cell isolation kit and cultured in RPMI-1640 medium further supplemented with 5% (v/v) NCTC-109 and 50 μM β-mercaptoethanol.

### Cell lines

The human embryonic kidney cell/Burkitt’s lymphoma fusion cell line HKB-11 was cultured in DMEM/F12 supplemented with 2 mM L-glutamine, 100 U/ml penicillin, 100 μg/ml streptomycin, 0.25 μg/ml amphotericin B and 10% FBS. WT, *Aire*^−/−^, *Acida*^−/−^ and *Ung*^−/−^ CH12 cells (generated in house) were cultured in RPMI-1640 medium further supplemented with 5% (v/v) NCTC-109 and 50 μM β-mercaptoethanol.

### Microbe strain

The *C. albicans* strain SC5314 was purchased from the ATCC, streak plates were made on YPD agar and incubated at 30°C overnight. Plates were then stored at 4°C. For experiments, single colonies were cultured in YPD broth at 30°C for 16 h with shaking at 220 rpm prior to induction of virulent pseudohyphae formation.

## METHOD DETAILS

### Human blood and tissue sample processing and cell isolation

Peripheral blood mononuclear cells (PBMCs) of APS-1 patients and healthy controls were purified using Ficoll-Paque Plus. Live (7AAD^−^ or Ghost Violet 510^−^) naive B cells (CD19^+^IgD^+^CD27^−^) and class-switched memory B cells (CD19^+^IgD^−^CD27^+^) were sorted from the PBMCs to a purity of ≥ 99% on a FACSAria II sorter (BD). PBMCs of anonymous healthy donors were isolated using a Histopaque-1077 gradient following the manufacturer’s instruction. Red blood cells (RBCs) were lysed using an ammonium-chloride-potassium (ACK) lysing buffer. Human tonsil and spleen tissues were minced into small pieces, meshed through 100 μm cell strainers, and pelleted at 600 g for 5 min at 4°C. Spleen cells were treated with an ACK buffer to remove erythrocytes and filtered through 40 μm cell strainers. Tonsil and spleen cells were then washed with phosphate-buffered saline (PBS). IgD^+^ B cells were purified from tonsil cells by magnetic-activated cell sorting (MACS) with a biotinylated goat F(ab’)_2_ anti-human IgD antibody and anti-biotin magnetic microbeads as previously reported (Chen et al., 2009). CD19^+^ B cells were similarly obtained using a biotinylated mouse anti-human CD19 (clone HIB19) antibody. The CD19^+^IgD^‒^ fraction was obtained by selecting for CD19^+^ cells from the IgD^‒^ fraction by MACS. Thymic cell suspensions were obtained by mincing human thymus tissues into small pieces and mechanically removing thymocytes followed by 2 rounds of digestion with 0.2% (w/v) Collagenase V and 0.1 mg/ml DNase I in Hank’s Balanced Salt Solution (HBSS) for 45 min at 37°C with shaking. The digested samples were filtered through 70 μm cell strainers and washed with PBS.

### Mouse blood and tissue cell isolation

Blood was collected from mice before euthanasia. spleen, inguinal lymph nodes, mesenteric lymph node and Peyer’s patches were collected after euthanasia. Adjacent fat and other tissues were removed before single cells suspensions were prepared, filtered through 100 μm cell strainer. RBCs from blood were removed by centrifugation on Histopaque 1077, and those in spleens were lysed using an ACK buffer. The cells were washed in PBS and counted before cell sorting, flow cytometry or purification by MACS. Resting B cells were isolated from the spleens of age- and sex-matched *Aire*^+/+^ or *Aire*^−/−^ littermates by MACS using a B Cell Isolate Kit. The purity of the isolated B cells ranged from 97‒99.6% as determined by flow cytometry based on CD19 and B220 staining.

### Mouse immunization

2.5×10^7^ purified *Aire*^+/+^ or *Aire*^−/−^ B cells were introduced via the tail vein into each recipient µMT littermate mouse. One day after the adoptive transfer, each recipient was intraperitoneally (i.p.) immunized with 1 dose of 100 μg NP_32_-KLH in Complete Freund’s Adjuvant and 3 doses of 100 μg NP_32_-KLH in Incomplete Freund’s Adjuvant once every week. Four days after the last immunization, mice were sacrificed, and blood and spleens were collected for ELISA, flow cytometry or cell sorting. In some experiments, mice were immunized 3 times, each with 200 μl of 2% sheep red blood cells in CFA for the initial immunization and IFA for the following immunizations. For **Figure S1H**, mice were immunized i.p. with 4×10^8^ SRBCs in PBS in a final volume of 200 μl/mouse with or without CFA.

### BM chimeras and B cell chimeras

BM cells were isolated from the tibias and femurs of age- and sex-matched CD45.1 *Aire*^+/+^ and CD45.2 *Aire*^−/−^ mice by flushing the BM cells with serum-free RPMI-1640 using 27G needles. Following RBC lysis, B220^+^ BM cells were depleted by MACS after labeling the BM cells with a biotinylated anti-mouse B220 antibody and anti-biotin microbeads. The B220^−^ cells from CD45.1 *Aire*^+/+^ and CD45.2 *Aire*^−/−^ BMs were mixed at the ratio of 1:1 and adoptively transferred via the tail vein into sex-matched primary μMT littermate recipients 1 d after these recipients received 10 Gy total body irradiation, with each recipient receiving a total of 1.5×10^7^ cells. BM reconstitution was allowed to proceed for 28 days. Splenic resting naive B cells were isolated from these primary μMT recipients by MACS using a B Cell Isolation Kit similarly as described above. Purified splenic B cells were counted, adjusted to a ratio of 1:1 and adoptively transferred via the tail vein into sex-matched secondary μMT littermate recipients, with each recipient receiving a total of 1.5×10^7^ cells. Mice were then immunized with NP_32_-KLH and adjuvants as described above.

### Discrimination of intravascular and tissue leukocytes

A published method was used to distinguish intravascular and tissue leukocytes in mice (Anderson et al., 2014). In this method, mice were intravenously administered with 6 μg biotinylated anti-CD45 antibody 2–3 minutes prior to anesthesia and cardiac perfusion with PBS, followed by immediate tissue harvest and digestion. Intravascular cells were subsequently stained with fluorochrome-conjugated streptavidin in combination with other cell surface markers before analysis by flow cytometry.

### Culture and stimulation or primary B cells

Peripheral blood IgD^+^CD27^‒^ naive B cells of healthy subjects or APS-1 patients were stimulated with 500 ng/ml soluble CD40L (sCD40L) and 100 ng/ml IL-4 or 100 ng/ml IFN-γ. Purified mouse splenic B cells were stimulated with 500 ng/ml sCD40L with or without 100 ng/ml IL-4, 100 ng/ml IL-21 or 25 μM CAPE. In some experiments, sCD40L was replaced with 5 µg/ml anti-CD40. To determine cell proliferation, the cells were labelled with carboxyfluorescein succinimidyl ester (CFSE) according to the manufacture’s protocol prior to culture. Alternatively, 10 μM 5-ethynyl-2’-deoxyuridine (EdU) was added to the culture medium for 6 hours before EdU incorporation was determined by flow cytometry using a Click-iT EdU Flow Cytometry Assay Kit according to the manufacturer’s protocol.

### Generation and validation of *Aire*^−/−^ CH12 cells

Several clones of *Aire*^−/−^ CH12 cells were generated by targeting the *Aire* gene using the CRISPR/Cas9 system as described previously (Ran et al., 2013). Single guide RNAs (sgRNA) targeting exon 1 or exon 3 of mouse *Aire* gene (GenBank AJ007715.1) were designed using the online tool at http://crispr.mit.edu. Sequences with the highest score for the respective region were selected. To express sgRNAs, pairs of oligonucleotides were synthesized and cloned into pSpCas9(BB)-2A-Puro plasmid as reported (Ran et al., 2013). The sgRNA expression plasmid was then transfected into CH12 cells using electroporation (square wave pulse at 200 V for 30 ms) in serum-free RPMI-1640 with 5 mM glutathione in a 4-mm cuvette. 24 hours after transfection, cells were resuspended in 125 ng/ml puromycin for 48 hours. After a brief expansion in puromycin-free media, single cell clones from transfected cells were screened for loss of the sgRNA targeting site using PCR. Clones with deletions in both alleles were identified by PCR. To determine the exact genomic modifications in each clone, the sgRNA-targeting sites were amplified with primer pairs spanning the targeting sites, and PCR products were sequenced directly using the respective forward primer. In addition, PCR products from clones 43 and 53 were cloned into the pGEM-T Easy vector and sequenced with T7 primer. All three mutant clones used were confirmed to harbor frameshift mutations on both alleles, resulting in termination shortly after the frameshift site. The potential off-target sites in the mouse genome for each guide were identified by the same online tool (http://crispr.mit.edu). Cas9 generally does not tolerate more than 3 mismatches (Hsu et al., 2013). All off-target sites in a potential gene-coding region with non-zero scores (up to 4 mismatches) were verified by sequencing to be intact. The lack of AIRE protein expression in these clones was finally confirmed by Western Blot.

### Plasmids

Full-length human *AIRE* cDNA sequence was cloned into pcDNA3.1(–) with tandem C-terminal Myc and 6-Histidine tag. Sequences coding various domains of AIRE were deleted using a Phusion Site-Directed Mutagenesis Kit using appropriate primers. Briefly, to delete a specific section of *AIRE* in the vector, a pair of outward primers was designed to amplify the remaining region together with the plasmid backbone. PCR product was then phosphorylated at 5’ end and ligated with Quick T4 ligase to recircularize it. Human AID was obtained by cloning full-length *AICDA* into pFLAG-CMV2 vector with an N-terminal FLAG tag. Domain-specific deletion mutants and G23S and E58A point mutants of AID were generated using the Phusion Site-Directed Mutagenesis kit using appropriate primers. The full-length *Egfp* sequence from pcDNA3-eGFP (from Dr. Thilo Hagen, National University of Singapore) was then cloned in frame to the C-terminus of mouse *Aire, AireΔNLS*, or *AireΔCARD* using blunt end ligation of PCR-amplified fragments.

### Transfection

10^6^ seeded HKB-11 cells were cultured to 70‒90% confluence and transfected with 4 μg plasmid DNA using Lipofectamine 3000 in Opti-MEM by following the manufacturer’s instruction. CH12 cells were transfected using the Amaxa cell line nucleofector kit V (Lonza). Cells were pelleted at 100 g for 10 min and resuspended in the electroporation buffer according to the manual. 10^6^ cells in 100 μl were mixed with 2 μg of target plasmids. The mixture was transferred to cuvettes for electroporation with Nucleofactor 2b (Lonza) using the program D-023. After electroporation, complete medium was immediately added to promote recovery. The electroporated cells were subsequently divided equally into 2 wells, with one well left unstimulated and the other stimulated with 1 μg/ml anti-CD40, 1 ng/ml TGF-β1 and 12.5 ng/ml IL-4 for 3 d.

### C. albicans culture

A single colony of *C. albicans* was cultured in YPD broth at 30 °C for 16 h with shaking at 220 rpm*. C. albicans* existed in the blastospore form after the 16 h culture. The concentration of the culture was quantitated using a hemocytometer. The culture was then diluted 1:10 with fresh YPD broth containing 10% (v/v) heat-inactivated FBS and grown at 37 °C for 3 h with shaking at 220 rpm. An aliquot of the culture was removed and examined under the microscope to ensure that 95% of blastospores switched to the virulent pseudohyphal form. The culture was pelleted by centrifugation at 4,000 rpm for 10 minutes, washed with PBS twice and resuspended in PBS at the concentration of 5×10^6^ CFU per 50 μl based on the quantitation of the culture 3 h ago. The pseudohyphae samples were used for either intradermal infection of mice or the preparation of heat-killed samples by treatment at 95 °C for 2 h followed by 3 rounds of sonication on ice at 30% maximum power for 5 seconds per round using a sonifier.

### Cutaneous *C. albicans* infection

5×10^7^ purified *Aire*^+/+^ or *Aire*^−/−^ B cells were introduced via the tail vein into each recipient µMT mouse littermate. Starting from the day of adoptive transfer, 5 doses each of 10^6^ CFU heat-killed *C. albicans* pseudohyphae were given intraperitoneally to each recipient mouse every 4 d. Four days after the last injection, mice were infected with 5×10^6^ CFU live *C. albicans* pseudohyphae in 50 μl PBS per spot at the deep dermis of the shaved dorsal region (Conti et al., 2014). The actual dose of infection was determined by immediately plating serial dilutions of the inoculum on YPD agar in triplicate, incubating the plates at 28 °C for 24 h and colony enumeration. The inoculum size per spot ranged between 3.8–12.3×10^6^ CFU in various experiments. Four days after the infection, blood was obtained prior to sacrificing the mice. The entire dermal injection site was excised for histological evaluation of fungal burden by Grocott’s methenamine silver (GMS) stain or by plating, or for determination of effector T cell response by flow cytometry. For GMS stain, the tissues were immediately fixed in 10% formalin overnight and embedded in paraffin before sectioning. For plating, each tissue was weighed, minced, grounded thoroughly and resuspended in sterile PBS. Serial dilutions of the suspensions were plated on YPD agar in triplicate and incubated at 28 °C for 24 h before colony enumeration. The fungal load was calculated as CFU per mg of tissue. For flow cytometry, the tissues were washed in FBS-free RPMI-1640 twice, minced and digested in FBS-free RPMI-1640 containing 0.7 mg/ml collagenase II, 2 mM EDTA and 25 mM HEPES at 37 °C for 1 h. The digested samples were passed through a 70 μm cell strainer, washed twice with RPMI-1640 containing 10% FBS, 2 mg/ml glutamine, 100 U/ml penicillin, 100 μg/ml streptomycin and 25 μg/ml amphotericin B. The samples were then cultured in this medium further supplemented with 500 ng/ml PMA, 500 ng/ml ionomycin and 1 μg/ml GolgiPlug at 37 °C for 5 h before being harvested for flow cytometric analysis.

### Immunoprecipitation

Cultured cells were harvested, washed with cold PBS twice and lysed with a CelLytic M buffer containing 1× protease inhibitor cocktail and 1× Halt phosphatase Inhibitor for 60 minutes on ice. The lysates were centrifuged at 18,000 g for 15 minutes at 4 °C. Protein concentration in the supernatants was determined by a BCA Protein Assay Kit. Equal amounts of lysate supernatants were used for immunoprecipitation with specific or isotype control antibodies using protein G magnetic beads (Cell Signaling 8740 or Thermo Fisher Scientific 88847) according to the manufacturers’ instructions.

### RNA extraction and quantitative real-time polymerase chain reaction

RNA was extracted from cells or tissues other than those from the APS-1 patients using TRIzol. cDNA synthesis was performed using the Superscript III first strand synthesis system in a thermocycler (Bio-Rad T100). qRT-PCR was performed with PowerSYBR Green Master Mix on a StepOnePlus instrument (Applied Biosystems) using pairs of sense and anti-sense primers targeting the genes of interest. For APS-1 patients’ peripheral blood IgD^+^CD27^‒^ B cells, following stimulation, the cells were washed and stored in RNAlater. Prior to RNA isolation, cells were pelleted at 5,000 g for 5 min and the RNAlater was removed. The cells were washed once with ice-cold PBS. RNA was isolated using the lysis and stop solutions in a Cells-to-C_T_ 1-step SYBR Green kit (Thermo Fisher Scientific A25601) and amplified using an iTaq Universal SYBR Green One-Step kit (Bio-Rad 172-5150) on a StepOnePlus instrument using pairs of sense and anti-sense primers targeting the genes of interest. The *ACTB* (*Actb*) gene was used as an internal control for normalization.

### Chromatin immunoprecipitation and quantitative real-time PCR

ChIP was performed using a ChIP assay kit based on the manufacturer’s instructions with slight modifications. Following 3 d of stimulation of 10^6^ CH12 cells as described above, formaldehyde was added to the culture to the final concentration of 1% and incubated for 10 minutes at 37 °C to crosslink chromatin. The cells were pelleted, washed twice in PBS, resuspended in an SDS lysis buffer (1% SDS, 10 mM EDTA, 50 mM Tris, pH 8.1) for 10 minutes on ice. DNA was sheared by 3 rounds of sonication on ice at 30% maximum power for 3 seconds per round using a sonifier (Thermo Fisher Scientific Q500). After centrifugation at 13,000rpm for 10 minutes, the supernatants were harvested, diluted 10-fold in a ChIP dilution buffer (0.01% SDS, 1.1% Triton X-100, 1.2 mM EDTA, 16.7 mM Tris-HCl, 167 mM NaCl, pH 8.1) containing protease inhibitors, and precleared with 50% protein A agarose/salmon sperm DNA slurry for 30 minutes at 4 °C with rotation. After setting aside an aliquot as input, An AID or control antibody was then added and incubated overnight at 4 °C with rotation, followed by the addition of 50% protein A agarose/salmon sperm DNA slurry for 1 h at 4 °C with rotation. The agarose was then pelleted and sequentially washed once with the low salt wash buffer, once with the high salt wash buffer, once with the LiCl wash buffer and twice with TE buffer, all of which were components of the kit. DNA in the bound chromatin was eluted from the beads using an elution buffer (1% SDS, 0.1 M NaHCO_3_, pH 8.0), reverse-crosslinked from proteins by incubation at 65 °C for 4 h in the presence of 200 mM NaCl, cleaned by 20 μg/ml RNase A treatment for 30 minutes at 37 °C followed by 40 μg/ml proteinase K treatment for 1 h at 45 °C, purified using phenol/chloroform extraction followed by ethanol precipitation with carrier glycogen according to the kit’s manual and resuspended in TE buffer for quantitative real-time PCR analysis using PowerSYBR Green Master Mix on a StepOnePlus instrument (Applied Biosystems). The fold enrichment of DNA was calculated using the ΔΔC_T_ method with control antibody-precipitated samples as an internal reference, and further compared among different CH12 cells and treatments.

### Protein extraction and Western Blot

1-2×10^6^ CH12 or primary B cells were pelleted and washed twice with ice-cold PBS, lysed with a pH 8.0 protein extraction buffer containing 20 mM Tris-HCl, 150 mM NaCl, 1% IGEPAL CA-630 (NP-40, Sigma-Aldrich I8896), 0.1% sodium dodecyl sulfate (SDS), 1 mM EDTA and protease and phosphatase inhibitor cocktail for 30 minutes on ice. Supernatants were collected after centrifugation, heated at 98 °C in SDS sample buffer with 4% β-mercaptoethanol for 5 minutes to denature proteins. Proteins were resolved in 4–20% Bis-Tris gels or 10% Tris-Glycine gels and transferred to 0.2 µm polyvinylidene fluoride (PVDF) membranes. The membranes were blocked with 5% (w/v) non-fat milk in Tris-buffered saline with Tween-20 for 30 minutes to 1 h, incubated with primary antibodies overnight at 4 °C and subsequently with secondary antibodies conjugated to HRP. Signals were visualized with clarity western-blot ECL substrate and exposed on autoradiograph films.

### Genomic uracil quantitation

The uracils in genomic DNA were quantified using AA6 as described previously (Wei et al., 2017; Wei et al., 2015). CH12 cells were harvested by centrifugation and lysed by incubating for 1 h at 37°C in Tris-EDTA buffer (TE) containing 2 μg/ml RNase A (Sigma-Aldrich R6513) and 0.5% SDS, followed by incubation with 100 μg/ml Proteinase K (Qiagen 19131) at 56°C for 3 h. DNA was isolated by phenol:chloroform (1:1) extraction and ethanol precipitation and dissolved in TE. The DNA was then digested with HaeIII (New England BioLabs R0108) and purified as described above. Digested genomic DNA was incubated with 10 mM O-Allylhydroxylamine hydrochloride (Sigma-Aldrich 05983) for 1 h at 37°C to block the pre-existing aldehydic lesions and the DNA was ethanol precipitated. The DNA was then treated with *E. coli* uracil DNA-glycosylase for 30 min at 37°C to excise uracils and create new abasic sites. The resulting abasic sites were labeled with 2 mM AA6 for 1 hour at 37°C and DNA was re-purified and dissolved in ddH_2_O. AA6-tagged DNA was labeled with 1.7 µM DBCO-Cy5 (Click Chemistry Tools A130) under Cu-free conditions by shaking the reaction mixture for 2 h at 37°C. Labeled DNA was purified using the DNA Clean and Concentrator kit (Zymo research D4014). Genomic DNA from *E. coli* CJ236 strain served as the uracil standard. WT and bisulfite-treated *E. coli* DNA served respectively as the negative and the positive controls. DBCO-Cy5-labeled DNA was spotted onto a positively charged zeta probe membrane (Bio-Rad 1620153) using a vacuum filtration apparatus and the membrane was scanned using a Typhoon 9210 phosphor imager (GE Healthcare). Cy5 fluorescence was analyzed using the ImageQuant software.

### Conventional flow cytometry

Cells were incubated with an Fc blocking reagent (Miltenyi Biotec 130-059-901 or Tonbo Biosciences 70-0161) and stained in PBS at 4°C with antibodies to various cell surface antigens. In the experiments that used NP_8_-FITC and NP_36_-PE to measure B cell affinity maturation, NP_8_-FITC was added to the cells and incubated for 15 min before NP_36_-PE was added. For the staining of intracellular molecules, cells were subsequently fixed and permeabilized using a CytoFix/CytoPerm kit or a Transcription Factor Buffer set. Isotype-matched control antibodies were used to define the baseline staining for the molecules of interest. Cells or beads stained with each fluorochrome were used to establish fluorescent compensation. 7-aminoactinomycin D (7-AAD, Tonbo Biosciences 13-6993-T500 or BD 559925) or Ghost Dye Violet 510 (GV510) was used to identify and exclude non-viable cells from the analysis. Events were acquired on an LSR II or LSR Fortessa flow cytometer (BD) and analyzed using FlowJo version 7 or 10 (Tree Star).

### Imaging flow cytometry

CD19^+^ B cells were purified from tonsillar cell suspensions by MACS with a biotinylated anti-CD19 antibody and anti-biotin microbeads. The cells were then incubated with an Fc blocking reagent and stained at 4°C with antibodies to surface IgD and CD38, fixed and permeabilized, and stained for AID and AIRE or with isotype control antibodies. Nuclei were counter stained with 4’,6-diamidine-2’-phenylindole dihydrochloride (DAPI). Tonsillar cells stained with each fluorochrome were used to establish fluorescent compensation. Cells were imaged on an ImageStream X Mark II imaging flow cytometer (Amnis) and data were analyzed using IDEAS 6.1 (Amnis).

### Immunofluorescence analysis

Frozen human tissues were stored at −80°C before 6-7 μm tissue sections were made using a cryostat (Leica CM1950). Sections were fixed with 4% paraformaldehyde, permeabilized in PBS containing 0.2% Triton X-100, incubated with Image-iT FX signal enhancer solution, blocked with PBS containing 1% BSA, 0.1% Triton X-100, 100 μg/ml human IgG (for human tissues only) and 10% serum from the source of the fluorochrome-conjugated antibodies, and stained with various combinations of primary antibodies against the molecules of interest, followed by appropriate fluorochrome-conjugated secondary antibodies. For Bcl-6 staining, the antibody used was not diluted and, instead, a large enough volume was used to cover the tissue section. For AIRE staining, a 1:25 dilution of the Miltenyi Biotec anti-AIRE-APC conjugated antibody was used. Nuclei were visualized with DAPI. Following washing, slides were mounted using a FluoroSave reagent and imaged on a confocal microscope (Zeiss LSM 780 or Leica TCS SP5). Pseudocolor images were processed using Photoshop CS6 (Adobe).

### ELISA

ELISA to determine NP-specific antibody affinity maturation was performed as previously described (Ballon et al., 2011) with minor modifications in the reagents. Briefly, each serum sample was titrated on both NP_29_-BSA- and NP_4_-BSA-coated microtiter plates. The ratio of binding to NP_4_-BSA and NP_29_-BSA is an indicator of relative Ig affinity maturation. Bound antibodies were detected using horseradish peroxidase (HRP)-conjugated goat-anti-mouse IgG1, IgG2b or IgG. The colorimetric reaction was terminated with the addition of an equal volume of 1 M H_2_SO_4_ and quantitated on a microplate reader (BioTek Epoch) at 450 nm. ELISA to determine IgG1 and IgA secretion by *ex vivo* stimulated mouse B cells was performed using a mouse IgG1 or IgA quantitation set. Anti-IL-17A, IL-17F and IL-22 autoantibodies in the mouse sera were measured using microtiters plates coated with 1 μg/ml recombinant murine IL-17A, IL-17F or IL-22. The plates were blocked with 10% BSA in PBS, washed, incubated with mouse serum samples, washed and then incubated with an alkaline phosphatase (ALP)-conjugated horse-anti-mouse IgG antibody (1:500). Following washing, the colorimetric reaction was developed using the BluePhos phosphatase substrate system and quantitated on a microplate reader (BioTek Epoch) at 620 nm. For the measurement of autoantibodies that block the binding of IL-17A, IL-17F and IL-22 to their receptors, mouse sera were incubated at 4°C overnight with HRP-conjugated IL-17A, IL-17F or IL-22 of the optimized concentrations, and the mixtures were added to microtiter plates coated with IL-17RA homodimer or IL-22Rα1 after the plates were blocked with 10% BSA in PBS. The plates were then incubated at room temperature for 1 hour and washed. The colorimetric reaction was developed and terminated with the addition of an equal volume of 1 M H_2_SO_4_ and quantitated on a microplate reader at 450 nm.

### IgHV repertoire and mutation analysis

Live (7-AAD^−^) unswitched (IgM^+^IgD^+^) or switched (IgM^−^IgD^−^) NP-specific B cells (CD19^+^B220^+^NP_36_^+^) in the spleens of immunized μMT recipients were sorted using a SONY SH800 cell sorter (SONY Biotechnology) and resuspended in RNAProtect solution. High-throughput IgHV repertoire profiling by RNA-Seq was performed at iRepertoire, Inc. (Huntsville, AL, USA). The raw sequences were processed and analyzed using the IMonitor 1.1.0 pipeline (Zhang et al., 2015). With this pipeline tool, each sequence was mapped to the *Mus musculus* germline V-D-J sequences (IMGT, http://www.imgt.org/vquest/refseqh.html) to identify the V, D and J gene segments and the CDRs, which were also used for clonal clustering. The sequences observed only once in a sample were filtered off to reduce the sequencing error. Subsequently, the sequences were normalized according to the number of cells in each sample. By comparing the sequence of each clone with the germline sequence, the mismatches of nucleotides were regarded as potential mutations. To eliminate PCR noise and sequencing errors, the first 25 bp of the sequences corresponding to the primer-binding site were excluded from the analysis, and the sequences were filtered if 3 successive mismatches were observed in them. Finally, the mutation rate for each IMGT position in the IgHV was calculated if the sequencing depth for that position was ≥10, and the frequency of each type of nucleotide substitution at these mutated positions were computed for each Ig isotype.

## QUANTIFICATION AND STATISTICAL ANALYSIS

### Statistical Analyses

All statistical analyses were performed using Excel or Prism. Graphs represent data from at least 2 independent experiments, each consisting of at least 3 biological replicates. The exact number of the biological replicates (*n*) in the presented data set are indicated in the figure legends. Results are expressed as mean ± SEM. Pair-wise statistical difference was assessed by the parametric *t*-test or the non-parametric Mann-Whitney *U* test depending on the distribution of the data, which is stated in the figure legends. Multiple group comparisons were performed using 1-way ANOVA with Tukey’s post hoc test. Differences were considered significant when *P* values were < 0.05. *P* values > 0.05 are either not shown, marked with NS (not significant), or indicated in the figure if they are close to 0.05. \**P* < 0.05, \*\**P* < 0.01, \*\*\**P* < 0.001.

## DATA AND SOFTWARE AVAILABILITY

### Software Availability

All software used for the data analysis in this study is commercially or freely available. The IMonitor software used for immune repertoire analysis has been published (Zhang et al., 2015) and can be downloaded at https://github.com/zhangwei2015/IMonitor.

## Notes

### Competing Interest Statement

The authors have declared no competing interest.

### Summary of Updates

This version has included additional supplemental data and detailed discussions.

